# Linked regulation of genome integrity and senescence-associated inflammation by p53

**DOI:** 10.1101/2023.11.20.567963

**Authors:** Karl N. Miller, Brightany Li, Hannah R. Pierce-Hoffman, Shreeya Patel, Xue Lei, Adarsh Rajesh, Marcos G. Teneche, Aaron P. Havas, Armin Gandhi, Carolina Cano Macip, Jun Lyu, Stella G. Victorelli, Seung-Hwa Woo, Anthony B. Lagnado, Michael A. LaPorta, Tianhui Liu, Nirmalya Dasgupta, Sha Li, Andrew Davis, Anatoly Korotkov, Erik Hultenius, Zichen Gao, Yoav Altman, Rebecca A. Porritt, Guillermina Garcia, Carolin Mogler, Andrei Seluanov, Vera Gorbunova, Susan M. Kaech, Xiao Tian, Zhixun Dou, Chongyi Chen, João F. Passos, Peter D. Adams

**Affiliations:** Cancer Genome and Epigenetics Program; Sanford Burnham Prebys MDI; La Jolla, CA, USA; Laboratory of Biochemistry and Molecular Biology; National Cancer Institute; National Institutes of Health; Bethesda, MD, USA; Department of Physiology and Biomedical Engineering; Mayo Clinic; Rochester, MN, USA; Robert and Arlene Kogod Center on Aging; Mayo Clinic; Rochester, MN, USA; NOMIS Center for Immunobiology and Microbial Pathogenesis; Salk Institute for Biological Studies; La Jolla, CA, USA; Center for Cancer Therapy; La Jolla Institute of Immunology; La Jolla, CA; Shared Resources; NCI-designated Cancer Center; Sanford Burnham Prebys MDI; La Jolla, CA, USA; Institute of Pathology; School of Medicine and Health; Technical University Munich (TUM); Munich, Germany; Departments of Biology and Medicine; University of Rochester; Rochester, NY, USA; Center for Regenerative Medicine, Department of Medicine; Massachusetts General Research Institute; Boston, MA, USA; Harvard Stem Cell Institute; Harvard University; Cambridge, MA, USA

## Abstract

Genomic instability and inflammation are distinct hallmarks of aging, but the connection between them is poorly understood. Understanding their interrelationship will help unravel new mechanisms and therapeutic targets of aging and age-associated diseases. Here we report a novel mechanism directly linking genomic instability and inflammation in senescent cells through a mitochondria-regulated molecular circuit driven by p53 and cytoplasmic chromatin fragments (CCF). We show, through activation or inactivation of p53 by genetic and pharmacologic approaches, that p53 suppresses CCF accumulation and the downstream inflammatory senescence-associated secretory phenotype (SASP), without affecting cell cycle arrest. p53 activation suppressed CCF formation by promoting DNA repair, and this is reflected in maintenance of genomic integrity, particularly in subtelomeric regions, as shown by single cell genome resequencing. Activation of p53 in aged mice by pharmacological inhibition of MDM2 reversed signatures of aging, including age- and senescence-associated transcriptomic signatures of inflammation and age-associated accumulation of monocytes and macrophages in liver. Remarkably, mitochondria in senescent cells suppressed p53 activity by promoting CCF formation and thereby restricting ATM-dependent nuclear DNA damage signaling. These data provide evidence for a mitochondria-regulated p53 signaling circuit in senescent cells that controls DNA repair, genome integrity, and senescence- and age-associated inflammation. This pathway is immunomodulatory in mice and a potential target for healthy aging interventions by small molecules already shown to activate p53.

## Introduction

Cellular senescence is a cell fate initiated by severe cellular stress and characterized by specific phenotypes, including stable cell cycle exit^1^ and development of a heterogeneous, pro-inflammatory senescence-associated secretory phenotype (SASP)^2^. The SASP contributes to chronic disease vulnerability and frailty in aged animals^3,4^, which has led to efforts in development of therapies targeting senescent cells or senescence phenotypes to promote healthy aging^5-8^. “Senolytic” approaches that target and remove senescent cells are one area of intensive investigation, but they are still in early stages of potential clinical translation^9^.

An alternative approach to modulate senescence function, a so-called “senomorphic” approach, targets the SASP specifically. Although senomorphic approaches show some promise for reducing chronic disease burden such as cancer *in vivo*^10-16^, the molecular regulation of SASP and its relevance to age-associated disease vulnerability are poorly understood. There are many known regulators of the SASP, including mTOR^10^, GATA4^17^, p38MAPK^18^ and p53^2^. Paradoxically, p53, a master inducer of senescence, has been reported to be a suppressor of SASP^2^. A molecular explanation of this paradox has been lacking.

The SASP is also induced by cytoplasmic DNA^19^ such as LINE1^20^ and mtDNA^21^. We have previously shown that the SASP is driven by a mitochondria-nucleus retrograde signaling pathway that promotes expulsion of cytoplasmic chromatin fragments (CCF) from the nucleus to the cytoplasm of senescent cells^22^. CCF are sensed by the cGAS/STING pathway, which activates the master transcriptional regulator of the SASP, NFkB^23-26^. However, the molecular regulation of CCF formation and the possibility of targeting this pathway in aged tissue *in vivo* remain unclear. Here, we identify a p53-regulated pathway that is subject to mitochondrial control, which suppresses CCF formation and the SASP, while also promoting DNA repair and genome integrity of senescent cells. We show that this pathway is pharmacologically targetable in cultured cells and mice, validating it as a target for senomorphic interventions and highlighting a novel function of p53 in safeguarding genome integrity of senescent cells.

## Results

### p53 suppresses CCF formation

The molecular regulation of CCF formation is poorly understood^27^. We have previously shown that the DNA repair protein 53BP1 is a suppressor of CCF formation and the SASP^22^. To determine the molecular role of 53BP1 in CCF formation, we developed a 53BP1 exogenous expression-based CCF suppression assay with modest overexpression of 53BP1 in irradiation-induced senescent IMR90 primary human fibroblasts. Western blot and qPCR for a panel of SASP factors and cell cycle regulators confirmed suppression of SASP without reverting the senescence-associated cell cycle arrest (Fig. S1A,B). This pattern of gene expression was confirmed by hierarchical clustering of differentially expressed genes by RNA-seq (Fig. S1D), where the primary cluster of genes suppressed by 53BP1 exogenous expression included genesets typically associated with the SASP, such as Legionellosis, Rheumatoid arthritis and IL-17 signaling (Fig. SE, Table S1). We next mutated known functional domains of 53BP1 and tested their ability to suppress CCF formation (Fig. S1F-G). Analysis of these 53BP1 mutants showed that deletion of the tandem BRCT domain significantly increased CCF formation compared to expression of wild-type 53BP1, with comparable levels of protein expression (Fig. 1A, Fig. S1F, G). These domains are involved in protein-protein interactions and are known to bind p53, especially p53 phosphorylated on serine 15 (pSer15-p53)^28,29^, suggesting a role for p53 in control of CCF formation. Confirming this interaction in senescent cells, pSer15-p53 coimmunoprecipitated with 53BP1 and vice versa (Fig. S1H,I). Consistent with the hypothesis that a 53BP1-p53 complex suppresses CCF formation, siRNA knockdown of p53, starting four days after senescence induction by irradiation, greatly increased CCF formation (Fig. 1B). This timepoint was chosen because at this time the cells have exited the cell cycle but have not produced many CCF (Fig. 1C). Conversely, activation of p53 after senescence induction by exogenous p53 expression, knockdown of MDM2, or by inhibition of MDM2 with two potent and selective clinical grade inhibitors, RG7388 and HDM201, suppressed CCF formation (Fig. 1B-D; Fig. S1J-L)). Confirming an on-target effect of the MDM2 inhibitor (MDM2i), suppression of CCF formation by RG7388 was dependent on p53 (Fig. 1B). siRNA-mediated knockdown of p53 and MDM2 was confirmed by western blot and immunofluorescence and revealed a strong negative correlation between p53 protein levels and CCF formation (Fig. S1M-P). MDM2i was titrated to the minimal dose required for SASP suppression as measured by expression of IL8, 12.5 nM RG7388 and 25 nM HDM201 (Fig. S2A-D). Low-dose RG7388 treatment decreased turnover of p53 in a cycloheximide-chase assay (Fig. S2E). This dose was not senolytic and did not revert cell cycle arrest as measured by EdU incorporation assay (Fig. 1E,F). Exogenous expression of p53 was also not obviously senolytic (Fig. S1K). Treatment of irradiation-induced senescent cells for an extended two-week timeframe with low dose RG7388 was also not senolytic and did not revert cell cycle arrest as measured by phosphorylation of cell cycle regulator RB (Fig. S1Q,R). RG7388 also suppressed CCF formation in an etoposide-induced model of senescence (Fig. S1S) but had less effect on CCF formation in oncogene- and replication-induced models of senescence (data not shown). Transcriptional profiling by bulk RNA-seq and qPCR of senescent cells treated with RG7388 or HDM201 showed a marked downregulation of inflammation-related gene ontologies associated with the SASP and expression of specific SASP genes, but did not affect expression of cell cycle-related genes (Fig. 1G, Fig. S2F-I, Table S1). This suppression was most marked among NFkB target genes^30^, and had a less marked effect on p53-^30^, p21-^31^, and p16-associated^31^ secretomes (Fig. S2I, Table S1). These data are consistent with a senomorphic effect of p53 activation, specific to the NFkB-dependent inflammatory phenotype of senescent cells, but not senolysis nor a reversion of the entire senescence phenotype. Altogether, these data show that p53 is a potent and selective suppressor of CCF and the SASP in DNA damage-driven models of senescence, and that this function is pharmacologically targetable for specific senomorphic effect by MDM2i.

**Figure 1:**
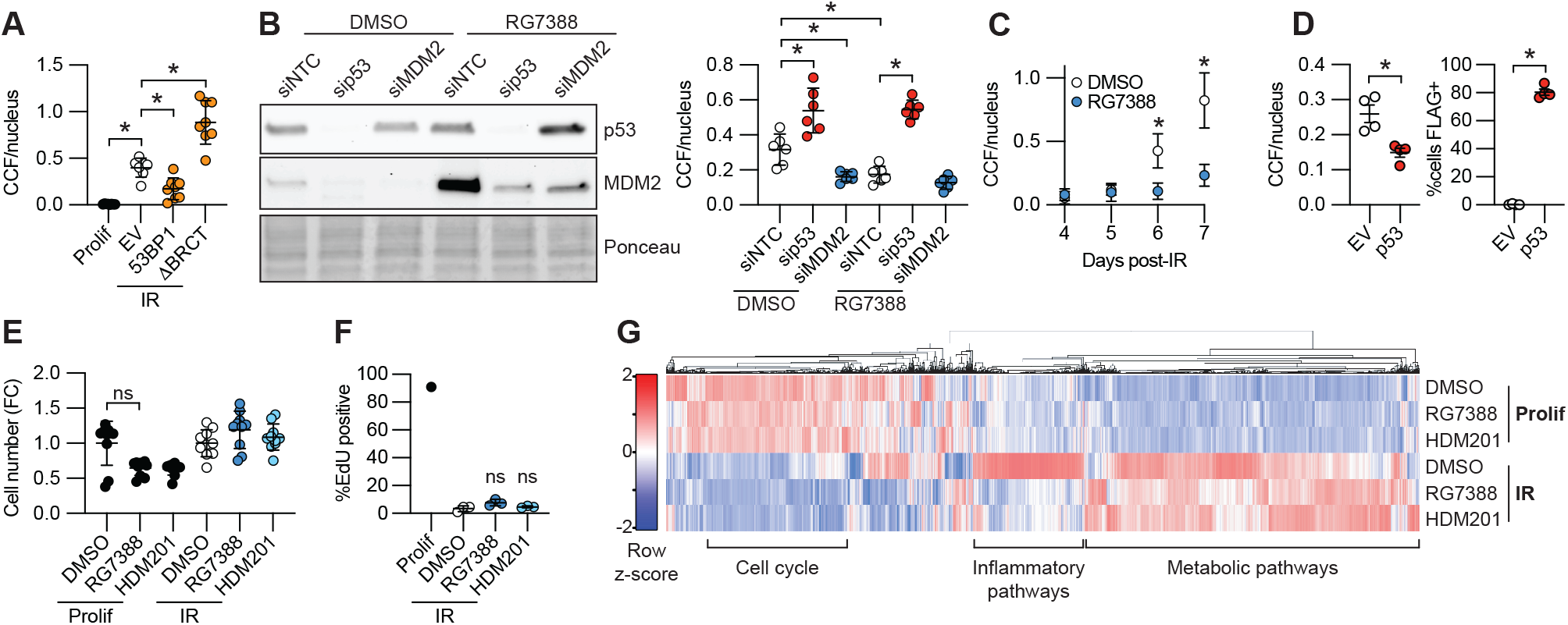
p53 suppresses CCF formation. A) CCF staining by IF in irradiation-induced senescent IMR90 human fibroblasts. Each value represents an individual well from a culture plate, representative of n=2 experiments. Y-axis represents total number of CCF normalized to total number of nuclei. B) Western blot and CCF staining by IF in irradiation-induced senescent IMR90 human fibroblasts using timeframe described in Fig. S1J, representative of n=2 experiments. C) CCF staining by IF at indicated day after irradiation, average of 3 experiments. D) CCF and FLAG staining by IF in irradiation-induced senescent IMR90 transduced with exogenous p53 or empty vector. Each value represents a separate infection in a single experiment, representative of n=3 experiments. E) Cell number as measured by number of nuclei, normalized to DMSO control for each group, representative of n=3 experiments. F) DNA replication as measured by EdU incorporation assay. Each dot represents a separate irradiation, representative of n=2 experiments. G) Differentially expressed genes by RNAseq, n=3 per group with summarized KEGG ontology for each major cluster. See Table S1 for detailed ontology. Data shown as means ±SD, asterisk(*) indicates p<0.05 by Student’s t-test. Prolif: proliferating control; EV: empty vector; NTC: non-targeting control; IR: ionizing radiation-induced senescence.

### p53 activation promotes DNA repair

The mechanism of CCF formation is thought to depend primarily on DNA damage and autophagy^22,32,33^. In senescent cells treated with MDM2i, we consistently observed a decrease in CCF, as well as nuclear γH2A.X foci and total γH2A.X protein levels by western blot (Fig. 2A-C). We also observed a decrease in nuclear γH2A.X foci with p53 exogenous expression (Fig. S1L) and with MDM2i treatment of etoposide-induced senescent cells (Fig. S1Q), as well as an increase in nuclear γH2A.X foci with p53 knockdown (Fig. S1O). These observations suggested that p53 activation may promote DNA repair in senescent cells. p53 is known to regulate DNA double-strand break (DSB) repair through both local activity directly at sites of DNA damage and transcriptional activation of DNA repair factors ^34,35^. Suggestive of p53 being a suppressor of CCF through a direct or local role in DNA repair^35^, pSer15-p53 colocalized with intranuclear γH2A.X in senescent cells as previously reported^36^, but was absent from CCF themselves (Fig. S3A), similar to the staining pattern of 53BP1 in senescent cells^32^. Also consistent with this model, suppression of CCF through activation of p53 with MDM2i or MDM2 knockdown increased colocalization of nuclear pSer15-p53 and γH2A.X foci at sites of intra-nuclear DNA damage in senescent cells, while p53 knockdown decreased colocalization (Fig. S3B-D). Most notably, across these treatments there was a positive correlation between intranuclear γH2A.X free of pSer15-pp53 and CCF, but an inverse correlation between pSer15-pp53 free of γH2A.X and CCF (Fig. S3D). However, the transcriptional role of p53 is also implicated in CCF suppression. Short-term treatment of senescent cells with RG7388 induced expression of p53 target genes, including several with known roles in DNA repair^37^, such as CDKN1A, XPC, BBC3, and PPM1D (Fig. 2D). CDKN1A (also known as p21) is thought to be required for efficient DNA repair, including repair of DSBs^38^ and has been shown to suppress DNA damage response (DDR)-associated toxicity in senescent cells^39^. Knockdown of p21 in senescent cells using a shortened timeframe to avoid cell death in the absence of p21 increased γH2A.X foci and CCF formation, even in the presence of MDM2i, showing that p53-mediated activation of DNA repair and suppression of CCF formation is dependent on its transcriptional target p21 (Fig. 2E). Consistent with a role for p53 activation in DNA repair, treatment of senescent cells with RG7388 decreased DNA double-strand break burden as measured by neutral comet assay (Fig. 2F). To confirm a role for p53 activation in DNA repair in senescent cells, we used an established NHEJ reporter system in I9A human fibroblasts^40-42^. This system uses doxycycline-inducible lentiviral expression of the rare-cutting endonuclease I-SceI to generate double-strand breaks around a silencing element within a GFP reporter cassette integrated into the I9A immortalized human fibroblast cell line, the NHEJ-dependent repair of which reconstitutes a functional GFP reporter. Lentiviral infection and doxycycline induction of I-SceI in I9A reporter cells four days after irradiation showed increased NHEJ efficiency with three days of RG7388 treatment (100 nM) compared to DMSO control by flow cytometry (Fig. 2G; Fig. S3F). RG7388 suppressed CCF formation and γH2A.X foci formation in irradiation-induced senescent I9A cells, similar to the effect we observed in IMR90 cells (Fig. S3E). We conclude that p53 activation promotes repair of DNA DSBs in senescent cells, a process that is very tightly linked to suppression of CCF.

**Figure 2:**
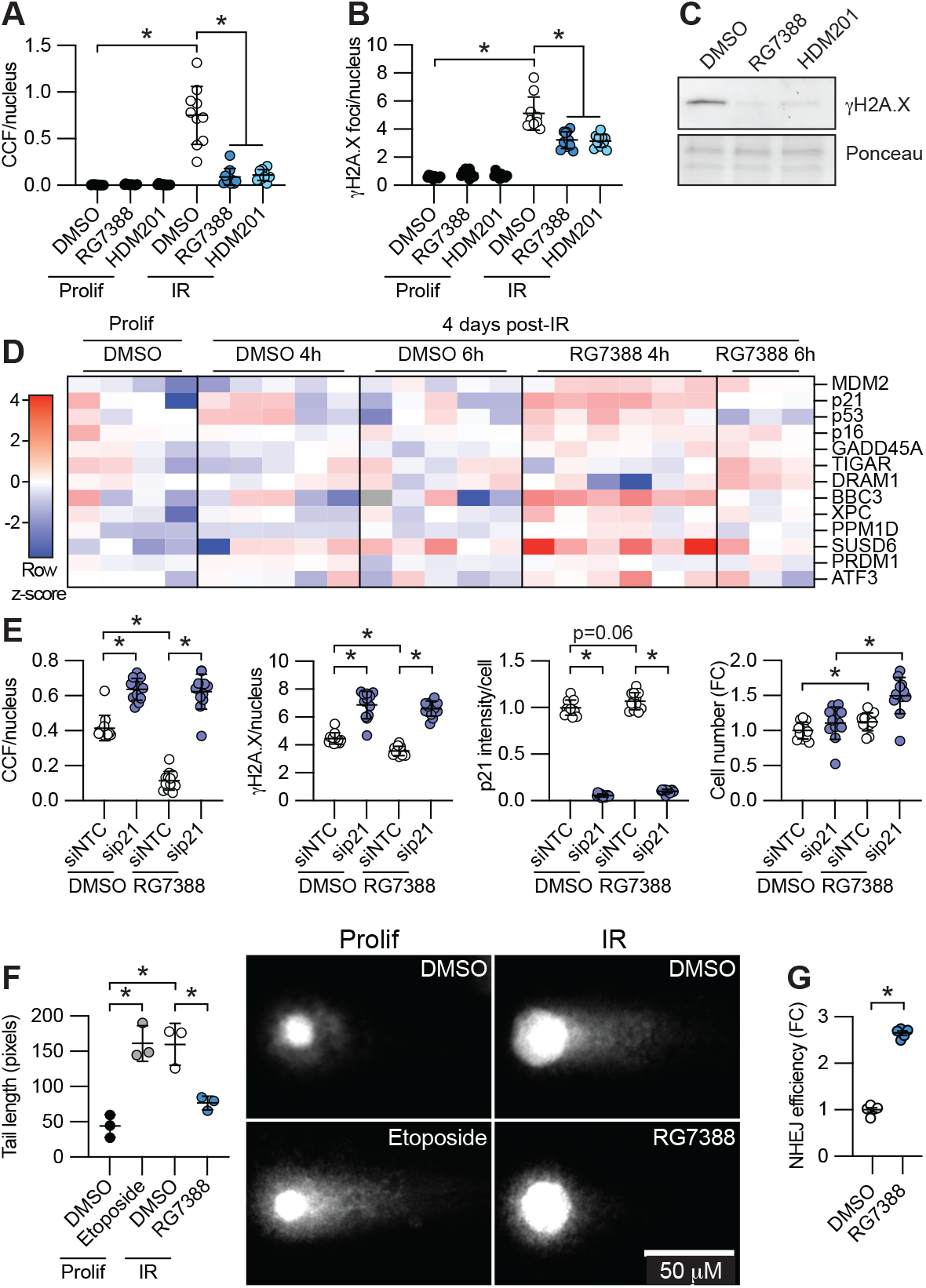
p53 activation promotes DNA repair. A) CCF formation and B) nuclear *γ*H2A.X foci number by IF in irradiation-induced senescent IMR90 human fibroblasts. Each value represents one well of a culture plate, from a representative experiment. C) WB of *γ*H2A.X in irradiation-induced senescent IMR90 cells from Fig. S1I, n=2 observations. D) qPCR analysis of p53 target genes in cells 4 days after irradiation, treated as indicated with RG7388 or DMSO control, n=3-6, representative of 3 experiments. E) IF quantitation of CCF, nuclear *γ*H2A.X, p21, and cell number quantified by number of nuclei in irradiation-induced senescent IMR90 cells. F) Neutral comet assay in irradiation-induced senescent IMR90 cells or proliferating control cells, representative of n=3 experiments. G) NHEJ reporter assay in irradiation-induced senescent I9A human fibroblasts, n=5 independent infections, representative of n=3 independent experiments. Data shown as means ±SD, asterisk(*) indicates p<0.05 by Student’s t-test. Prolif: proliferating control; NTC: non-targeting control; IR: ionizing radiation-induced senescence; NHEJ: non-homologous end joining.

### p53 preserves genome integrity in senescent cells

We reasoned that p53’s ability to promote DNA repair in senescent cells should underpin a role in preservation of genome integrity of senescent cells. To test this, we irradiated IMR90 cells and assessed genome copy number by single-nucleus genome resequencing after full establishment of senescence. Cells were treated with RG7388 five minutes after IR. In the absence of RG7388, single cell genome library preparation with a linear amplification-based approach (LIANTI)^43^ showed large amplifications and deletions across the genomes of irradiated senescent cells with substantial heterogeneity, relative to normal proliferating cells that were diploid with few copy number variations (Fig. S4A). This was confirmed by a second PCR-based approach (Fig. 3A, Fig. S4B). Remarkably, activation of p53 with RG7388 preserved genome integrity, as marked by diploid genome content with very few copy number variations in 5/6 cells sequenced (Fig. 3A,B, Fig. S4B). Interestingly, we observed that these deletions rarely included pericentromeric sequences and tended to occur towards the telomeric ends of chromosomes (Fig. 3C). Consistent with this observation, we found that less than 10% of CCF contain the centromeric marker CENPA (Fig. 3D,F). However, detection of telomeric sequences in senescent cells by FISH confirmed that approximately 80% of CCF contain telomeric DNA (Fig. 3E,F). We conclude that p53 activation after acute genotoxic stress preserves genome integrity, presumably due to enhanced DNA repair, and this is tightly linked to suppression of telomeric DNA-containing CCF in senescent cells.

**Figure 3:**
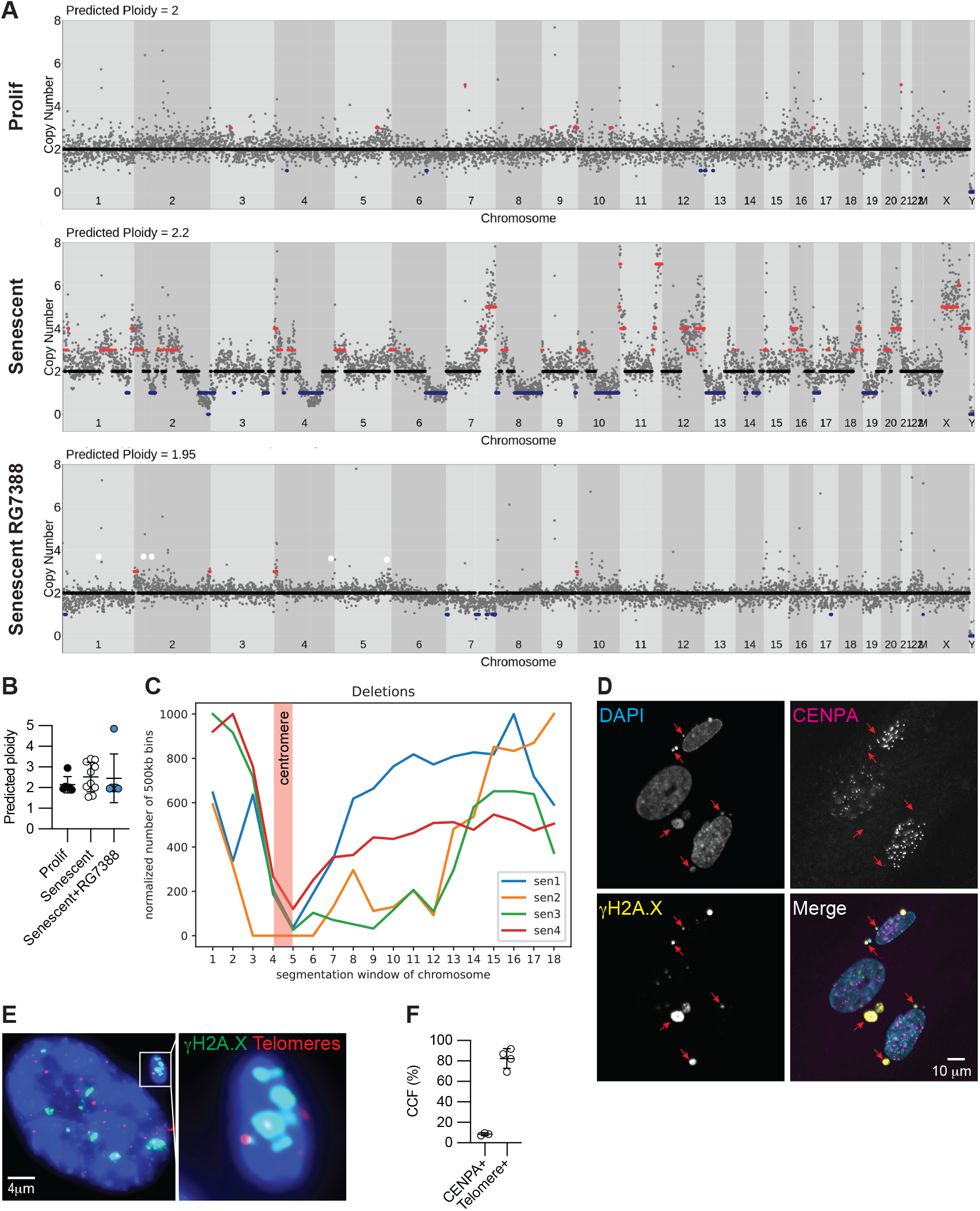
p53 preserves genome integrity in senescent cells. A) Representative single nucleus whole-genome plots showing copy number variations and B) predicted ploidy in irradiation-induced senescent IMR90 human fibroblasts, n=6, 10, 6 nuclei for prolif, Senescent, and Senescent RG7388 groups respectively. C) Histogram of deletions, with each line an aggregation of all chromosomes per cell, n=4 senescent cells. Each chromosome is divided into 18 equal bins, where bins 1 and 18 are subtelomeric and bins 4-5 are pericentromeric. D) IF for CENPA in irradiation-induced senescent IMR90 cells, with arrows marking CCF, representative of n=3 experiments and E) ImmunoFISH for telomeres and *γ*H2A.X, representative of n=4 independent experiments. F). Quantitation of D and E, where each CENPA+ marker represents n=3 separate irradiations from a representative experiment and each telomere+ marker represents n=4 independent experiments. Data shown as means ±SD, asterisk(*) indicates p<0.05 by Student’s t-test. Prolif: proliferating control.

### MDM2i is senomorphic in vivo

MDM2 inhibitors have been extensively developed as candidate cancer therapeutics and are generally considered highly specific *in vivo*^44,45^. To determine whether p53 activation can suppress SASP in vivo, we treated naturally aged mice with the clinical grade MDM2i, HDM201. Although HDM201 has been previously shown to have specific, on-target activity in mouse and rat tumor models^46,47^, little is known about its activity in normal tissue, especially aged tissue. To test the effect of HDM201 on senescence phenotypes *in vivo*, we treated aged male and female mice for 14 days. This regimen was not associated with changes in weight, whole blood counts, or indices of liver pathology in either sex, suggesting that the treatment was well-tolerated (Fig. S5A-C). We focused our molecular analysis on liver, which is known to accumulate senescent cells and inflammation-associated pathologies with age^48^. Protein levels of p53 and downstream target p21 measured by western blot were significantly increased in HDM201-treated mice compared to vehicle, confirming on-target engagement of the drug (Fig. 4A,B). Interestingly, bulk RNA-seq of whole liver tissue showed extensive changes in gene expression in female mice treated with HDM201 compared to vehicle control, but few changes in gene expression were observed in male mice (Table S2), revealing a sex-dimorphic effect at this dose. In female mice, targeted analysis of established p53 target genes^49^ showed a trend for increased expression of 45/103 genes (Fig. S5D), and statistically significant increased expression for 12 of these genes (Table S2). Assessment of liver senescent cell burden in female mice by telomere-associated DDR foci (TAF) staining showed an increase in TAF with age compared to young vehicle controls (5 month), but no change with HDM201 treatment compared to vehicle control (Fig. 4C), consistent with the selective SASP-suppressive non-senolytic effect of MDM2i we observed in cultured cells. Unbiased analysis showed 4912 genes differentially expressed in old vehicle-treated female mice compared to young vehicle female controls. Among these 4912 genes, 824 were differentially expressed with HDM201 treatment, of which the vast majority (776) showed reversal of the age-associated gene expression change (Fig. S5E). Remarkably, this was also apparent across all 4912 genes differentially expressed with age (Fig. 4D), suggesting that HDM201 treatment reverses an age-associated transcriptional signature. Genes downregulated with age and upregulated with drug treatment were associated with sterol and fatty acid metabolic pathways, while genes upregulated with age and downregulated with drug treatment were associated with immune and inflammation pathways (Fig. 4E, Table S2). Ingenuity pathway analysis (IPA) of all differentially expressed genes confirmed this pattern, in which expression of genes associated with upstream regulators of SASP and inflammation (for example TNF, IFNG, TFGB1, and STAT1) increased as a function of age, but decreased as a function of HDM201 treatment (Fig. 4F,G, Table S2). HDM201 treatment did not significantly decrease age-associated fibrosis or adiposity (Fig. S5F,G). However, consistent with a SASP-suppressive effect of HDM201 treatment, HDM201 reversed age-associated accumulation of macrophages and dendritic cells in liver of aged mice (Fig. 2H, Fig. S8). These data show that p53 activation in aged female mice does not alter senescent cell burden, but reverses an age-associated transcriptional signature and immune cell composition, supporting an anti-inflammatory, senomorphic, and immunomodulatory activity of MDM2i *in vivo*.

**Figure 4:**
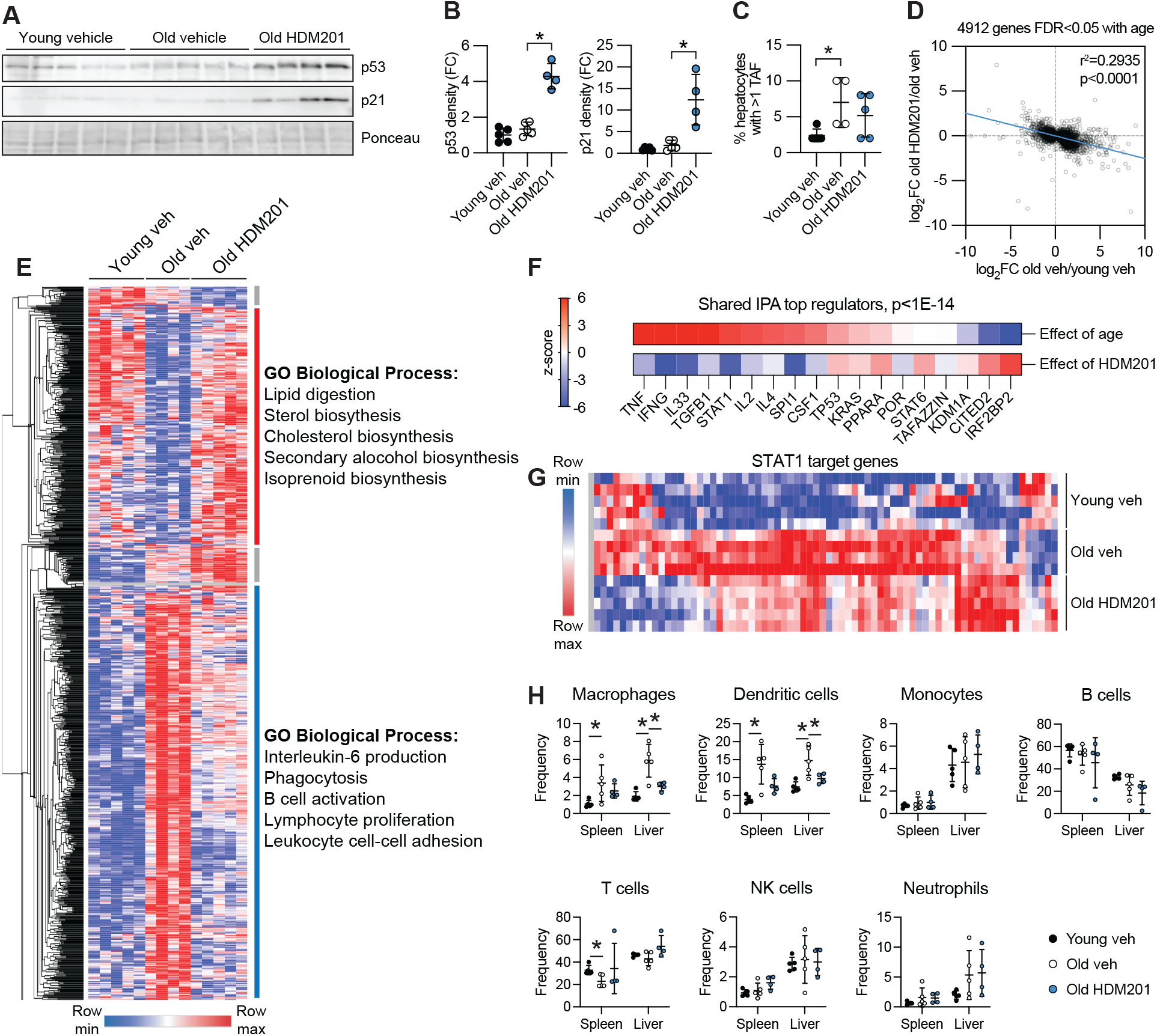
MDM2i is senomorphic in vivo. A) WB of p53 and p21 in mouse liver and B) quantitation, n=4-5 per group. C) TAF assay in female mice showing the percentage of hepatocytes with greater than 1 TAF, n=4-5 per group. D) Correlation between change in gene expression as a function of age and change in gene expression as a function of HDM201 treatment, among 4912 DE genes with age, n=4-5 per group. E) Heatmap combining 776 reversed DE genes, where the change in expression of a gene is opposed by HDM201 treatment, and 58 genes not reversed, with top 5 GO biological process terms for major hierarchical clusters (see also Fig. S5E), n=4-5 per group. F) Ingenuity pathway analysis of DE genes showing top upstream regulators common between old vehicle vs. young vehicle (effect of age) and old HDM201 vs. old vehicle (effect of HDM201) comparisons, using a cutoff of p<1E-14. G) Representative IPA target gene heatmap of STAT1. H) Flow cytometry analysis of immune cell frequencies isolated from spleen and liver, n=4-5. Data shown as means ±SD, asterisk(*) indicates p<0.05 by Mann-Whitney U test or (H) one-way ANNOVA. TAF: telomere-associated DNA damage response foci; Veh: vehicle control; DE: differentially expressed.

### Mitochondria suppress p53 activity in senescence

We next sought to understand upstream regulators of p53 in the context of CCF formation. We have previously shown that mitochondrial stress drives CCF formation in senescent cells, and ablation of mitochondria by Parkin-mediated forced mitophagy^50^ in senescent cells suppresses CCF formation^22^ and SASP^22,51^. Thus, mitochondria and p53 are seemingly antagonistic regulators of CCF in senescence. To test the relationship between mitochondrial stress and p53, we irradiated IMR90 cells, then ablated mitochondria by Parkin-mediated forced mitophagy. After cell cycle exit and mitochondrial ablation, we knocked down p53 by siRNA (Fig. S6A,B). Consistent with our previous work, ablation of mitochondria suppressed CCF formation, increased intranuclear γH2A.X foci number^22^, and suppressed expression of SASP marker IL8 (Fig. 5A). However, although knockdown of p53 alone increased CCF formation and expression of SASP, cells with both ablation of mitochondria and knockdown of p53 had very few CCF and showed greatly increased and altered γH2A.X staining, with marked redistribution of γH2A.X staining throughout the nucleus, compared to cells with mitochondria treated with non-targeting siRNA (Fig. 5B, Fig. S6B). This result shows that ablation of mitochondria is dominant to inactivation of p53 in terms of CCF formation and SASP but not in terms of intranuclear γH2A.X accumulation, suggesting that mitochondria are required for CCF formation and expression of the SASP at a step downstream of p53-mediated DNA repair.

**Figure 5:**
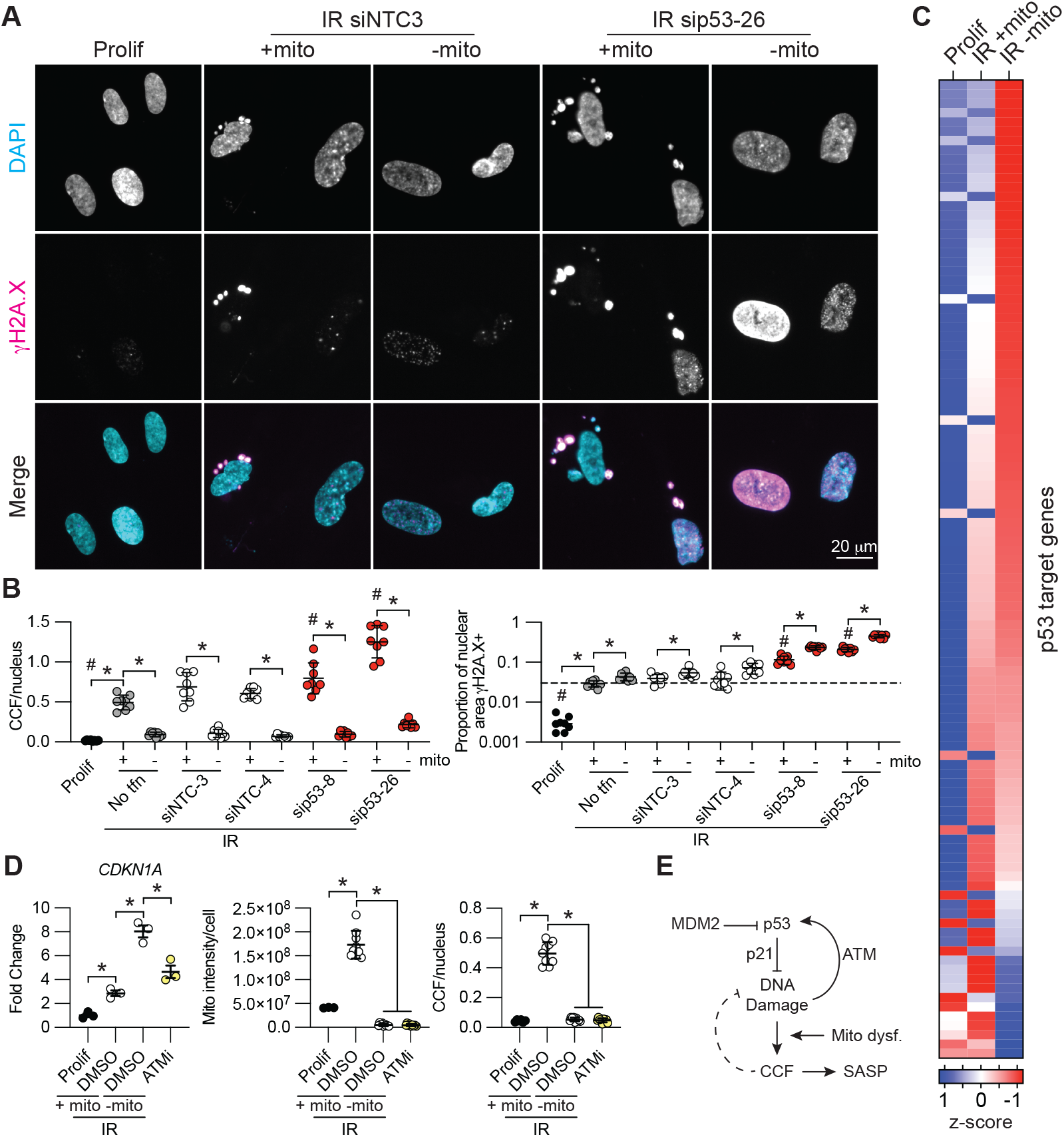
Mitochondria suppress p53 activity in senescence. A) IF representative images and B) quantitation of CCF and proportion of the nucleus staining positive for *γ*H2A.X in irradiation-induced senescent Parkin-overexpressing IMR90 human fibroblasts. See Fig. S6A for experimental timeline. Each value represents an individual well from a culture plate, from a representative experiment, n=3. Numbering indicates individual siRNA sequences. C) p53 target gene expression by RNAseq from ref^22^, average of n=3 per group shown. D) qPCR of CDKN1A and corresponding IF quantitation of mitochondria content and CCF in irradiation-induced senescent Parkin-over-expressing IMR90 human fibroblasts. See Fig. S6E for experimental timeline. E) Model. Data shown as means ±SD, asterisk(*) indicates p<0.05 by t-test, (#) in panel B indicates p<0.05 vs siNTC-4. Prolif: proliferating control; NTC: non-targeting control; No tfn: no transfection control; IR: ionizing radiation-induced senescence.

Because CCF remove damaged DNA from the nucleus, a driver of p53 activation, we further reasoned that mitochondrial stress-associated CCF formation might indirectly regulate p53 activity by decreasing nuclear DNA damage burden. Specifically, we hypothesized that mitochondrial ablation in senescent cells would activate p53 by increasing nuclear retention of damaged DNA. Indeed, ablation of mitochondria in senescent cells greatly induced expression of p53 target genes in two different primary fibroblast strains (Fig. 5C, Fig. S6C,D) and was accompanied by increased intranuclear γH2A.X and pS15-p53 (Fig. 5B, Fig. S6B), consistent with the idea that p53 activation results from elevated DNA damage and the DNA damage response pathway. To test this directly, we inhibited the kinase ATM, which activates p53 in response to DNA damage by phosphorylation of serine 15^52,53^. Pharmacological inhibition of ATM in senescent cells blocked transcriptional activation of p21 in response to ablation of mitochondria, suggesting that mitochondrial control of p53 activity is, at least in part, ATM-dependent (Fig. 5D, Fig. S6E). These data show that mitochondria in senescent cells suppress p53 activity, in part through an ATM-dependent feedback pathway.

## Discussion

These data provide evidence for a mitochondria-regulated p53-CCF circuit in senescent cells (Fig. 5E). This circuit controls DNA repair, genome integrity and SASP. p53 suppresses CCF formation and the SASP tightly linked to DNA repair, a pathway which is subject to feedback regulation by mitochondria. We show that activation of this pathway preserves genome integrity in senescent cells. The p53-CCF circuit is a potential target for anti-inflammatory and genome-stabilizing healthy aging interventions.

We note some limitations to this study. First, we acknowledge that some effects of MDM2i may extend to p53-independent pathways. However, in our cell culture model, we show that MDM2i suppression of CCF formation is dependent on p53 (Fig. 1B), likely due to stabilization of p53 protein levels (Fig. S2E). Second, we acknowledge that the senomorphic effects of MDM2i are primarily observed in DNA damage-driven irradiation- and etoposide-induced models of senescence, with much weaker effects on CCF formation in oncogene-induced senescence and replicative senescence (data not shown). This is perhaps not surprising, given the DNA repair-associated mechanism by which p53 suppresses CCF formation (Fig. 2). However, elevated DNA damage is an established marker of senescence *in vivo*^54,55^, and we show evidence that this pathway is relevant to senescence-associated phenotypes and immune function in aged mice (Fig. 4), suggesting that this p53-CCF pathway is relevant to the natural aging process.

By single nucleus genome re-sequencing, we observed large deletions on the order of tens of millions of kilobases that tended to occur at the telomeric ends of chromosomes. CCF contained telomeres but not centromeres, suggesting the simple hypothesis that double-strand breaks are more likely to generate CCF if they are close to the end of a chromosome. Consistent with this, evidence from the literature shows that specific generation of DNA damage at telomeres is sufficient for CCF formation^56^. Because centromeres are 2-3 orders of magnitude larger than telomeres^57^, we might expect that CCF would contain more centromeric DNA by random chance. However, we do not observe this, which is consistent with a non-random genomic origin of CCF. However, we cannot formally exclude an alternate model, that the mechanism of chromatin loss from the ends of chromosomes is instead dependent on preferential retention of centromeres, which are known to be distended in senescence^58^. p53 is known as the “guardian of the genome”, in part by suppressing genome instability in precancerous cells^35^. Although many mechanisms are implicated, recent work has shown that p53 loss in the course of cancer evolution leads to deterministic loss of genome integrity on an individual cell basis^59,60^, perhaps consistent with our observation of a cell-intrinsic role for p53 in promoting genome stability in irradiation-induced senescence.

It has previously been reported that p53 suppresses the SASP^2,61^, consistent with the observation that p53 activity declines in senescence after cell cycle exit, while SASP increases^62^—but the mechanism of SASP suppression by p53 has been unclear. We show that p53 suppresses formation of CCF, which are known to activate the SASP through a cGAS-STING pathway ^23-26^. Additionally, this finding integrates our previous observations that dysfunctional mitochondria in senescent cells drive CCF formation and SASP ^22,51^. Altogether, we propose a model in which p53 preserves genome integrity on a single-cell basis by promoting repair of DNA DSBs. In senescent cells, this pathway is suppressed by mitochondrial dysfunction, which instead drives resolution of DNA damage by cytosolic expulsion of the damaged DNA as CCF. This expulsion appears to indirectly downregulate p53 and to require a second mitochondria-driven pathway independent of p53, perhaps related to autophagy^33^.

The role of p53 in aging is unclear—genetic manipulation of the p53 pathway is associated with either longevity or accelerated aging, depending on context^63^. The MDM2i UBX0101 has been explored as a senolytic in the context of osteoarthritis^64^, although a phase 2 clinical trial failed to demonstrate efficacy in humans^65^. Recently, the MDM2i BI01 was shown to have senolytic activity in aged mouse muscle^66^. However, in our in vitro models, we observe only SASP-suppressive senomorphic activity, not senolytic activity. We show the MDM2i HDM201 is potentially senomorphic in mouse liver. These effects were observed primarily in female mice, perhaps due to the treatment regimen used, which was established to minimize toxicity observed primarily in female mice at higher doses (data not shown). The suppression of the inflammation-associated gene expression signature and the immunomodulation we observe is consistent with a growing literature on the anti-inflammatory effects of MDM2i in vivo^67^. A better understanding of MDM2 inhibitors in experimental aging models could be a useful tool in the pursuit of interventions to promote healthy aging in humans.

## Methods

### Animals

This study was approved by the Institutional Animal Care and Use Committee at Sanford Burnham Prebys MDI. C57BL6 mice were purchased from Jackson Labs and Charles River and group housed under specific pathogen free conditions with ad libitum access to water and food (Teklad 2018). Mice were treated with 10 mg/kg HDM201 (Novartis) suspended in phosphate-buffered methylcellulose according to the manufacturer’s instructions (0.5% methylcellulose in phosphate buffer, pH6.8, 50mM) by daily oral gavage in the morning. Mice were monitored daily and weighed every few days during treatment. After 14 days of treatment, mice were euthanized by CO2 asphyxiation in the morning, and tissues were fixed in 10% neutral buffered formalin (Epredia 9400-1) or flash frozen in liquid nitrogen, either whole or in OCT (Tissue-Tek 4583). Whole blood was counted using a hematology analyzer (Abaxis VetScan HM5).

### Cell culture, senescence induction, and drug treatments

IMR90 primary human fibroblasts were grown at 37°C, 3.5% O2, 5% CO2, in Dulbecco’s modified Eagle’s medium (Gibco 10313-121) with 10% FBS (Corning 35-0-11-CV), 1% penicillin/streptomycin (Gibco 15140-122) and 2 mM glutamine (Gibco 25030-081). Cells were checked routinely for mycoplasma contamination. IR senescence was induced by 20 Gray x-ray irradiation of 20-30% confluent cells. Cells were split after returning to confluence in 3 days, then treated with MDM2 inhibitors, RG7388 (Selleckchem) and HDM201 (Novartis), starting day 4 after irradiation or day 5 after irradiation in experiments that also included siRNA treatment, unless otherwise noted. Unless otherwise indicated, cells were collected 10 or 11 days after irradiation. For longer-term treatments, drugs were replaced by media change every 3 days. RG7388 was used at 12.5 nM, or 100 nM (Fig. 1B; Fig. S1S; Fig. 2G; Fig. S3E,F), and HDM201 was used at 25 nM. IMR90 cells exogenously expressing Parkin and the method for mitochondria ablation are previously described (Vizioli) and 10 μM KU55933 was used for ATM inhibition experiments. For etoposide-induced senescence, 80% confluent IMR90 cells were treated with 50 mM etoposide for 48 hours, RG7388 was added on day 4 after initiation of senescence, and cells were collected on day 7. 293T cells were purchased from ATCC. I9A human fibroblast cells^40^ were a gift from Dr. Vera Gorbunova.

### Plasmids

pLV-hPGK-HA-53BP1-puro and pLV-EF1A-FLAG-p53-puro constructs were generated by Vectorbuilder. Mutations to 53BP1 are described in Fig. S1. pLV-TetOne-eBFP2-I-SceI-Puro was a gift from Dr. Vera Gorbunova.

### Lentivirus infection

For 53BP1 exogenous expression, lentivirus was generated from 293T cells transfected with expression vector, VSVG envelope vector, and pMD2.G packaging vector in lipofectamine 2000 (Invitrogen 52887). Virus was collected over 3 days, then concentrated by ultracentrifugation. For p53 exogenous expression, concentrated lentivirus was generated by the SBP viral vector core facility using a proprietary protocol. Concentrated virus was titrated to the minimum amount required for >90% cell viability after puromycin selection (1 μg/mL for 72 hours). Infections were overnight (16 hours), in the presence of 8 μg/mL polybrene (Millipore TR-1003-G).

### siRNA transfection

Senescent cells were transfected 4 days after irradiation with 100 nM siRNA (Dharmacon siGENOME) in 0.8% Dharmafect reagent (Dharmacon) according to manufacturer’s recommendations. Experiments used a pool of four siRNAs per gene unless otherwise noted. Cells were changed into regular media 18-20 hours later, except sip21 experiments.

### Antibodies

The following primary antibodies were used: 53BP1 (Cell Signaling Technology Cat#4937, RRID:AB_10694558), ATM (D2E2) (Cell Signaling Technology Cat#2873, RRID:AB_2062659), CENPA (Thermo Fisher Scientific Cat#MA1-20832, RRID:AB_2078763), Cyclin A (Santa Cruz Biotechnology Cat#sc-271682, RRID:AB_10709300), ANTI-FLAG M2 (Sigma-Aldrich Cat#F3165-2MG, RRID:AB_259529), Phospho-Histone H2A.X (Ser139) (Millipore Cat#05-636, RRID:AB_309864, Active Motif Cat#39117, RRID:AB_2793161), HA-probe (F-7) (Santa Cruz Biotechnology Cat#sc-7392, RRID:AB_627809), IgG control (Vector Laboratories Cat#I-1000, RRID:AB_2336355, Vector Laboratories Cat#I-2000, RRID:AB_2336354), IL8 (Abcam Cat#ab18672, RRID:AB_444617), MDM2 (Cell Signaling Technology Cat#51541, RRID:AB_2936381), p21 (Santa Cruz Biotechnology Cat#sc-817, RRID:AB_628072), p53 (Santa Cruz Biotechnology Cat#sc-126, RRID:AB_628082, Leica Biosystems Cat#NCL-p53-CM5p, RRID:AB_563933), Phospho-p53 (Ser15) (Cell Signaling Technology Cat#9284S, RRID:AB_331464), Phospho-ATM (Ser1981) (Abcam Cat#ab81292, RRID:AB_1640207), TOMM20 (Abcam Cat#ab56783, RRID:AB_945896). The following secondary antibodies were used: Goat anti-Mouse IgG, IgM (H+L) HRP (Thermo Fisher Scientific Cat#31446, RRID:AB_228318), Goat anti-Rabbit IgG, (H+L) HRP (Millipore Cat#AP307P, RRID:AB_92641), Goat anti-Mouse IgG (H+L), Alexa Fluor^TM^ 594 (Thermo Fisher Scientific Cat#A11032, RRID:AB_2534091), Goat anti-Rabbit IgG (H+L), Alexa Fluor^TM^ 488 (Thermo Fisher Scientific Cat#A11008, RRID:AB_143165). Antibodies used for flow cytometry are listed in Table S3.

### Western blot

For cells, blotting was done as previously described^68^. Briefly, cells were lysed in modified RIPA buffer (50 mM Tris-Cl pH 7.5, 0.25% sodium deoxycholate, 150 mM NaCl, 10 mM EDTA, 0.1% SDS, 1% Igepal, 1x dual protease and phosphatase inhibitor (Thermo 1861281)) and lysates were cleared by >20000g centrifugation. Protein was quantified by BCA assay (Pierce 23225), mixed with sample buffer, run on precast gels (Biorad), and transferred either with a Turbo semidry system (Biorad), or for 53BP1 experiments wet transfer in tris-glycine buffer (Biorad) with 5% methanol. Membranes were blocked in milk and imaged by ECL (Thermo 34095, Biorad Chemidoc). For mouse liver tissue, samples were lysed in modified RIPA buffer using a Bertin Technologies Precellys tissue disruptor.

### Immunoprecipitation

Protein G Dynabead-antibody complexes were prepared as previously described^22^. Cells were washed 4 times in PBS, scraped in EBC500 (50 mM Tris-Cl pH 8, 500 mM NaCl, 0.5% Igepal, 2.5 mM MgCl2) with benzonase (250 U/mL), and lysed by rotating 30 minutes at 4°C. Protein was quantified by BCA assay. Immunoprecipitations were run overnight at 4°C, washed 7 times in NETN (20 mM Tris-Cl pH 8, 100 mM NaCl, 1 mM EDTA, 0.5% Igepal), and eluted in sample buffer.

### Immunofluorescence

Cells were plated on PhenoPlate^TM^ 96-well microplates (PerkinElmer), stained as described previously^22^, and imaged on a Nikon T2 microscope with automated image capture. Images were analyzed in NIS Elements using rolling ball background subtraction, thresholding, size exclusion, and automated partitioning to identify features. Measurements indicate total number of features divided by total number of nuclei, except in Figure 5, where total mitochondrial or nuclear γH2A.X staining area is divided by total number of nuclei.

### TAF staining and Immuno FISH

Staining was done as previously described^21^.

### Comet assay

This assay was carried out following manufacturer’s instruction for the neutral comet SCGE assay (Enzo, #ADI-900-166). Slides were placed flat in the dark at 4C in the dark for 30 minutes. Slides were immersed in pre-chilled lysis solution (2.5M NaCl, 100mM EDTA pH10, 10mM Tris Base, 1% sodium lauryl sarcosinate, 1% Triton x-100, Catalog No.4250-050-01) for 45 minutes. Electrophoresis was performed in TAE buffer (40mM Tris base, 20mM acetic acid, 1mM EDTA disodium salt dihydrate) at 30V for 15 minutes at room temperature. Automated comet analysis was performed using an open-source tool in ImageJ (Gyori et al 2014).

### qPCR

Cells were lysed in Trizol (Ambion 15596026) and RNA was isolated using either a commercial kit (53BP1 and MDM2i RNAseq experiments; Direct-zol RNA Miniprep Kits, Zymo Research Cat#R2050) or with chloroform according to the manufacturer’s protocol. RNA was converted to cDNA (RevertAid Reverse Transcriptase, Thermo Fisher Scientific Cat#EP0441, Ribolock RNase Inhibitor, Thermo Fisher Scientific Cat#EO0381, 5x reaction buffer for RT, Thermo Fisher) and gene expression quantified by a standard SYBR-based approach (PowerUp^TM^ SYBR^TM^ Green Master Mix for qPCR, Applied Biosystems Cat#A25741, QuanStudio^TM^ 6 Flex Real-Time PCR System, 384-well, Applied Biosystems Cat#4485691).

### RNA-seq

RNA was quantified by bioanalyzer and library preps were made by the SBP genomics core. Sequencing was done by the SBP genomics core or at UCSD Institute for Genomic Medicine. For analysis, raw fastq files were aligned to hg19 (53BP1 OE RNAseq) or hg38 (MDM2i in cell culture RNAseq), or mm10 (MDM2i in mice RNAseq) using STAR^69^ 2-pass pipeline. Reads were filtered, sorted and indexed by SAMtools^70^. FPKM were generated using CuffLinks^71^ for downstream visualization. Genome tracks (bigWig files) were obtained by Deeptools^72^. Raw read counts were obtained by HTSeq^73^ for differential analysis. Differentially expressed genes were obtained by DESeq2^74^. KEGG gene ontology was run using WebGestalt^75^, gene lists were compared using Venny2.1(https://bioinfogp.cnb.csic.es/tools/venny/index.html), and heatmaps were generated using Morpheus (https://software.broadinstitute.org/Morpheus).

### Single cell genome resequencing

Cells were trypsinized, washed in PBS, and resuspended at 20x cell pellet volume in cold nuclear isolation buffer A with digitonin (50 nM HEPES pH 7.3, 150 mM NaCl, 1x dual protease and phosphatase inhibitor, 25 μg/mL digitonin) by pipetting. The suspensions were rotated at 4°C for 30 minutes, then centrifuged at 500g for 5 minutes at 4°C. Nuclear pellets were washed twice with cold NIB-250 buffer (250 mM sucrose, 15mM Tris-Cl pH 7.5, 60 mM KCl, 15 mM NaCl, 5 mM MgCl2, 1 mM CaCl2) and resuspended in sorting buffer (DPBS with 2% FBS, 0.5 mM spermidine, 500 ng/mL DAPI). Nuclei were sorted by FACSAriaII with a 100um nozzle (see also Fig. S7) into individual strip tubes, and libraries were generated by LIANTI^43^ or PicoPLEX Gold (Takara Bio). Libraries were sequenced by NOVAseq (Illumina). For analysis, the first 14 bases of both R1 and R2 reads were trimmed using CutAdapt^76^. Trimmed reads were then aligned to hg38 using Bowtie2^77^. The sam files were transferred to bam files, then sorted and indexed using SAMtools^70^. Duplicates were removed using picard tools (https://broadinstitute.github.io/picard/) MarkDuplicates function. Copy number variation profiles were obtained via Ginkgo^78^.

### NHEJ reporter assay

I9A cells were irradiated at 20 Gy, split 1:2 three days after irradiation, then on the next day infected with lentivirus containing I-SceI and treated simultaneously with 1 μg/mL doxycycline and 100 nM RG7388 or DMSO. 16 hours later, virus was removed and treatments were refreshed. Cells were collected on day seven after irradiation and analyzed by flow cytometry using a BD LSRFortessa Cell Analyzer.

### Cycloheximide chase assay

IMR90 cells were pretreated for 30 minutes with 12.5 nM RG7388 or DMSO, then treated in reverse with 100 μM cycloheximide for 1 or 3 hours. The 0 hour control was treated with DMSO for 3 hours.

### Mouse immune cell profiling by flow cytometry

Tissues were dissected and placed on ice-cold RPMI supplemented with 10% FBS. Single-cell suspensions of immune cells from the liver were obtained by mechanical disaggregation through a 70µm cell strainer (VWR) and washed through with 10% FBS in RPMI. Liver samples were spun at 60 r.c.f. and 4ºC for 2 min with no brake to pellet hepatocytes before percoll (Cytiva) centrifugation. The supernatant was collected, spun at 420 r.c.f. and 4ºC for 4 min. The pellet was resuspended with 40% Percoll (Cytiva) in HBSS to further remove debris and hepatocytes. The isolated immune cells from the liver went through red blood cell lysis with ACK buffer (KD Medical) before counting cells on a hematocytometer. Splenocytes were isolated by passing cells through a 70µm cell strained followed by red blood cell lysis with ACK buffer before being transferred to a 96-well U-bottom plate and resuspended in fluorescence-activated cell sorting (FACS) buffer (2% FBS in 1X PBS). Viability staining was performed using LIVE/DEAD fixable red stain (1 in 1000 in FACS buffer, Invitrogen) for 15 min at room temperature. Suspensions were then pelleted and resuspended in anti-CD16/32 antibodies (1:500, BioLegend) to block non-specific binding of Fc receptors. Cells were incubated with the indicated surface antibodies for 30 min at 4ºC. a FoxP3 transcription factor staining kit (eBioscience) was used for intracellular staining. Antibodies against intracellular proteins were diluted in 1X permeabilization buffer and added for 45 min at 4ºC. For cytokine staining, cells were stimulated with PMA (final concentration of 1 µg/mL) and ionomycin (Iono, Cell Signaling; final concentration of 1 µg/mL) for 4h at 37ºC in the presence of brefeldin A (GolgiPlug, BD Biosciences; final concentration of 1 µg/mL) to block cytokine export from the golgi apparatus. 2% paraformaldehyde (PFA) was used to fix the cells after staining. Cells were resuspended in 100µL 1X PBS and run on the FACSymphony A3 5-laser flow cytometer (BD Biosciences). Data were analyzed using FlowJo (v.10, BD Biosciences).

### Histology

Formalin-fixed liver tissue was paraffin embedded, and sectioned using standard approaches. H&E, Picosirius Red, and oil red-O staining was done by the SBP histology core using standard approaches and analyzed using Python 3.6.10. H&E images were scored by a trained pathologist (C.M.) for age-associated liver pathology. For Picosirius Red, images were processed by applying a median filter to each RGB channel using a disk of size 2, then converted to HSV to isolate red hues with predefined thresholds. Noise was reduced by removing small objects, and the Picosirius Red-stained areas were quantified as a ratio of stained area or intensity to the total non-white area. For oil red-O, images were processed using the OpenCV library for red droplet isolation via HSV color segmentation, followed by a 2×2 pixel morphological opening to refine droplet boundaries. Droplet count, size, and total area were quantified using skimage.measure.

### Statistical analysis

Statistical significance for routine assays was calculated in Graphpad Prism, using p<0.05 as a threshold. For cell culture experiments, pairwise comparisons were done by two-sided Student’s t-test assuming unequal variance between groups. Simple linear regressions were calculated in Graphpad Prism. For animal experiments, pairwise comparisons were done using a Mann-Whitney U test, or by one-way ANOVA (Fig. 4H).

## Acknowledgements

We would like to thank Adriana Charbono for technical support with mouse experiments. This work was supported by K99 AG073450 and F32 AG066459 (to KNM); CIRM EDUC4-12813 (to MGT); the Glenn Foundation for Medical Research Postdoctoral Fellowship PD19131 and the AFAR Reboot Fund (to ND); P30CA03199 (YA); R00AG068303 (to XT); R01AG082785 (to ZD); the Intramural Research Program of the National Institutes of Health, National Cancer Institute (1ZIABC011884) (to JL and CC); R37046320 (to AS), P01AG047200 and R01AG027237 (to VG); R01AG68048, UG3CA268103 and R01AG82708 (to JFP) and P01 AG031862 and R01 AG071861 (to PDA).

## Author contributions

Conceptualization: KNM, BL, HRPH, SP, CCM, TL, ND, ZD, PDA. Methodology: AR, MGT, ND, ABL, SL, AD, JL, AK, AS, VG, SMK, XT, YA, RP, GG, CC. Data generation and analysis: KNM, BL, HRPH, XL, APH, SP, AG, CCM, MGT, AR, SGV, MAL, SW, ABL, AD, EH, ZG, YA, CM, XT. Writing: KNM, BL, ZD, PDA. Funding: KNM, PDA. Supervision: CC, JFP, PDA.

## Materials & Correspondence

Correspondence and requests for materials should be addressed to KNM and PDA.

## Data availability

All sequencing data are deposited in GEO (data deposition in progress). All other data are available upon reasonable request.

## Figure legends

**Figure S1: Related to Figure 1**. A) WB and B) qPCR of 53BP1 and senescence markers in irradiation-induced senescent IMR90 human fibroblasts ectopically expressing 53BP1 or empty vector. C) RNA-seq principal component analysis, D) hierarchical clustering of differentially expressed genes, and E) KEGG ontology of cluster indicated by dashed box. F) 53BP1 protein map with major domains targeted by mutagenesis. G) CCF formation by IF and WB of 53BP1 mutant protein expression in irradiation-induced senescent IMR90 human fibroblasts ectopically expressing wild-type or mutant 53BP1 or empty vector. H,I) IP of pS15-p53 or 53BP1 in irradiation-induced senescent IMR90 human fibroblasts, n=1 experiment each. J) Schematic of siRNA experiments. K) Cell number and L) nuclear γH2A.X foci by IF related to Fig. 1D, representative of n=3 experiments. M-P) Deconvolution of siRNA pools used in Fig. 1B, assessing CCF, p53, and nuclear γH2AX by IF, n=1 experiment. Q) Irradiation-induced senescent IMR90 fibroblasts treated with 12.5 nM RG7388 for 14 days, showing markers of senescence and p53 activation by western blot and R) cell number, CCF formation, nuclear γH2A.X, and cleaved caspase 3 intensity by IF, representative of n=2 experiments. S) Etoposide-induced senescent IMR90 fibroblasts showing CCF formation, nuclear γH2A.X, and cell number, representative of n=2 experiments. Data shown as means ±SD, asterisk(*) indicates p<0.05 by Student’s t-test. Prolif: proliferating control; IR: ionizing radiation-induced senescence; ETO: etoposide; wt: wild-type; EV: empty vector; OE: overexpression; en: endogenous transcript; ex: exogenous transcript; NTC: non-targeting control; IR: ionizing radiation-induced senescence; NT: n-terminal domain; UDR: ubiquitin-dependent region; NLS: nuclear localization sequence; BRCT: Brca1 C-terminal sequence motif.

**Figure S2: Related to Figure 1**. A-D) MDM2i dose-response curves in irradiation-induced senescent IMR90 human fibroblasts by IL8 WB, n=1 experiment each. E) Cycloheximde chase assay in IMR90 cells 4 days after irradiation, measuring p53 protein level by WB and quantitation as fold change vs. 0 h for each group, representative of n=2 experiments. F) qPCR validation and G) RNA-seq principal component analysis for MDM2i treatment related to Fig. 1G. H) Venn diagram of DE genes. I) Left to right: NFkB, p53, p21, and p16-associated secretomes. Data shown as means ±SD, asterisk(*) indicates p<0.05 by Student’s t-test. P: proliferating control; IR: ionizing irradiation-induced senescence; CHX: cycloheximide.

**Figure S3: Related to Figure 2**. A) Representative IF image of γH2A.X and pS15-p53 colocalization in irradiation-induced senescent IMR90 human fibroblasts and B) IF quantitation of colocalization between pS15-p53 and γH2A.X, representative of n=2 experiments. C) Quantitation of γH2A.X foci not colocalized with pS15-p53, and D) correlations to CCF formation of (left) γH2A.X foci not colocalized with pS15-p53 foci and (right) pS15-p53 foci not colocalized with γH2A.X foci, from same dataset as Fig. 1B. E) Irradiation-induced senescent I9A human fibroblasts, showing CCF formation and nuclear γH2A.X foci by IF, representative of n=2 experiments. F) Representative raw data related to Fig. 2G. Data shown as means ±SD, asterisk(*) indicates p<0.05 by Student’s t-test. Prolif: proliferating control; NTC: non-targeting control.

**Figure S4: Related to Figure 3**. Whole genome copy number variation plots for all cells sequenced by A) LIANTI or B) PicoPLEX Gold approaches.

**Figure S5: Related to Figure 4**. A) Body weight and change in weight over the course of treatment as a percentage of baseline animal weight, n=4-10 per group. B) Whole blood measures and white blood cell composition, n=4-10. C) Liver pathology indices scored by a trained pathologist, n=4-10. D) p53 target gene expression in female mouse liver by RNA-seq, n=4-5 per group. E) Comparison of genes differentially expressed with age and HDM201 in liver by bulk RNA-seq, n=4-5 per group. F) Picosirius red staining in female mice, n=9-10. G) Oil red-O staining, n=4-5. Data shown as means ±SD, asterisk(*) indicates p<0.05 by Mann-Whitney U test. V: vehicle; H: HDM201; Y: young; O: old; F: female; M: male.

**Figure S6: Related to Figure 5**. A) Timeline for mitochondrial ablation experiments in Fig. 5 A,B. B) WB validation of mitochondrial ablation and p53 knockdown related to Fig. 5A,B. C) p53 target genes with mitochondrial ablation in MRC5 fibroblasts, from ref^51^. D) Heatmap related to Fig. 5C, showing additional controls. E) Timeline for mitochondrial ablation experiment with ATMi treatment in Fig. 5D. IR: ionizing radiation-induced senescence.

**Figure S7: Gating of cell sorting for single-nucleus genome resequencing**. A) Proliferating, B) Senescent, C) Senescent + MDM2i. Red marks correspond to the individual sorted nuclei from Fig. S4B.

**Figure S8: Flow cytometry gating for immune profiling related to Figure 4**. A) Representative gating strategy for identifying immune cells mouse liver. B) Representative flow cytometry plots showing the frequency of macrophages (F4/80+CD64+) and C) dendritic cells (CD11c+) isolated from liver and spleen. The plots in B and C are gated on CD45+CD11b+Ly6C-cells.

**Table S1: RNAseq analysis from cell culture experiments**.

**Table S2: RNAseq analysis from mouse experiments**.

**Table S3: List of antibodies used for immune cell profiling**.

