## Supplementary material for "Linked regulation of genome integrity and senescence-associated inflammation by p53": Suplemental Figures

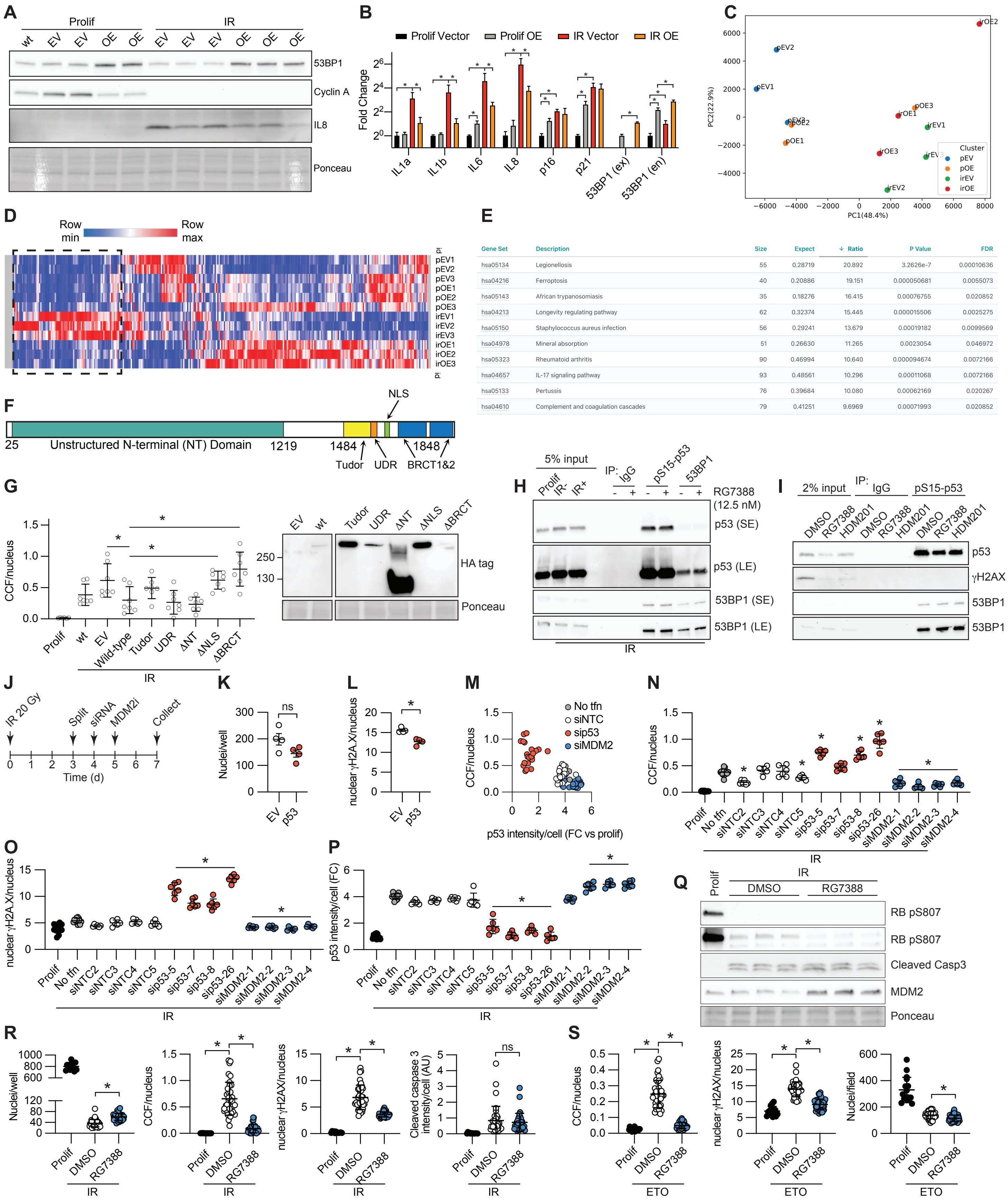
A
Prolif
IR
wt EV EV OE OE EV EV EV OE OE OE
53BP1
Cyclin A
IL8
Ponceau
B
Prolif Vector
Prolif OE
IR Vector
IR OE
Fold Change
IL1a IL1b IL6 IL8 p16 p21 53BP1 (ex) 53BP1 (en)
C
PC2(22.9%)
PC1(48.4%)
Cluster
pEV
pOE
irEV
irOE
D
Row min
Row max
pEV1 pEV2 pEV3 pOE1 pOE2 pOE3 irEV1 irEV2 irEV3 irOE1 irOE2 irOE3
E
Gene Set | Description | Size | Expect | Ratio | P Value | FDR
hsa05134 | Legionellosis | 55 | 0.28719 | 20.892 | 3.2626e-7 | 0.00010636
hsa04216 | Ferroptosis | 40 | 0.20886 | 19.151 | 0.000050681 | 0.0055073
hsa05143 | African trypanosomiasis | 35 | 0.18276 | 16.415 | 0.00076755 | 0.020852
hsa04213 | Longevity regulating pathway | 62 | 0.32374 | 15.445 | 0.000015506 | 0.0025275
hsa05150 | Staphylococcus aureus infection | 56 | 0.29241 | 13.679 | 0.00019182 | 0.0099569
hsa04978 | Mineral absorption | 51 | 0.26630 | 11.265 | 0.0023054 | 0.046972
hsa05323 | Rheumatoid arthritis | 90 | 0.46994 | 10.640 | 0.000094674 | 0.0072166
hsa04657 | IL-17 signaling pathway | 93 | 0.48561 | 10.296 | 0.00011068 | 0.0072166
hsa05133 | Pertussis | 76 | 0.39684 | 10.080 | 0.00062169 | 0.020267
hsa04610 | Complement and coagulation cascades | 79 | 0.41251 | 9.6969 | 0.00071993 | 0.020852
F
NLS
25 Unstructured N-terminal (NT) Domain 1219 1484 1848
Tudor UDR BRCT1&2
G
CCF/nucleus
Prolif wt EV Wild-type Tudor UDR ΔNT ΔNLS ΔBRCT
IR
EV wt Tudor UDR ΔNT ΔNLS ΔBRCT
250
130
HA tag
Ponceau
H
5% input
IP: IgG pS15-p53 53BP1
Prolif IR- IR+
RG7388 (12.5 nM)
p53 (SE)
p53 (LE)
53BP1 (SE)
53BP1 (LE)
IR
I
2% input
IP: IgG pS15-p53
DMSO RG7388 HDM201 DMSO RG7388 HDM201 DMSO RG7388 HDM201
p53
γH2AX
53BP1 (SE)
53BP1 (LE)
J
IR 20 Gy
Split siRNA MDM2i Collect
Time (d)
K
Nuclei/well
EV p53
ns
L
nuclear γH2A.X/nucleus
EV p53
M
No tfn
siNTC
sip53
siMDM2
CCF/nucleus
p53 intensity/cell (FC vs prolif)
N
CCF/nucleus
Prolif No tfn siNTC2 siNTC3 siNTC4 siNTC5 sip53-5 sip53-7 sip53-8 sip53-26 siMDM2-1 siMDM2-2 siMDM2-3 siMDM2-4
IR
O
nuclear γH2A.X/nucleus
IR
P
p53 intensity/cell (FC)
IR
Q
Prolif
IR
DMSO RG7388
RB pS807
RB pS807
Cleaved Casp3
MDM2
Ponceau
R
Nuclei/well
CCF/nucleus
nuclear γH2AX/nucleus
Cleaved caspase 3 intensity/cell (AU)
Prolif DMSO RG7388
IR
S
CCF/nucleus
nuclear γH2AX/nucleus
Nuclei/field
Prolif DMSO RG7388
ETO

**Figure S1: Related to Figure 1.** A) WB and B) qPCR of 53BP1 and senescence markers in irradiation-induced senescent IMR90 human fibroblasts ectopically expressing 53BP1 or empty vector. C) RNA-seq principal component analysis, D) hierarchical clustering of differentially expressed genes, and E) KEGG ontology of cluster indicated by dashed box. F) 53BP1 protein map with major domains targeted by mutagenesis. G) CCF formation by IF and WB of 53BP1 mutant protein expression in irradiation-induced senescent IMR90 human fibroblasts ectopically expressing wild-type or mutant 53BP1 or empty vector. H,I) IP of pS15-p53 or 53BP1 in irradiation-induced senescent IMR90 human fibroblasts, n=1 experiment each. J) Schematic of siRNA experiments. K) Cell number and L) nuclear $\gamma$H2A.X foci by IF related to Fig.1D, representative of n=3 experiments. M-P) Deconvolution of siRNA pools used in Fig.1B, assessing CCF, p53, and nuclear $\gamma$H2AX by IF, n=1 experiment. Q) Irradiation-induced senescent IMR90 fibroblasts treated with 12.5 nM RG7388 for 14 days, showing markers of senescence and p53 activation by western blot and R) cell number, CCF formation, nuclear $\gamma$H2A.X, and cleaved caspase 3 intensity by IF, representative of n=2 experiments. S) Etoposide-induced senescent IMR90 fibroblasts showing CCF formation, nuclear $\gamma$H2A.X, and cell number, representative of n=2 experiments. Data shown as means ±SD, asterisk(*) indicates $p<0.05$ by Student's t-test. Prolif: proliferating control; IR: ionizing radiation-induced senescence; ETO: etoposide; wt: wild-type; EV: empty vector; OE: overexpression; en: endogenous transcript; ex: exogenous transcript; NTC: non-targeting control; IR: ionizing radiation-induced senescence; NT: n-terminal domain; UDR: ubiquitin-dependent region; NLS: nuclear localization sequence; BRCT: Brca1 C-terminal sequence motif.

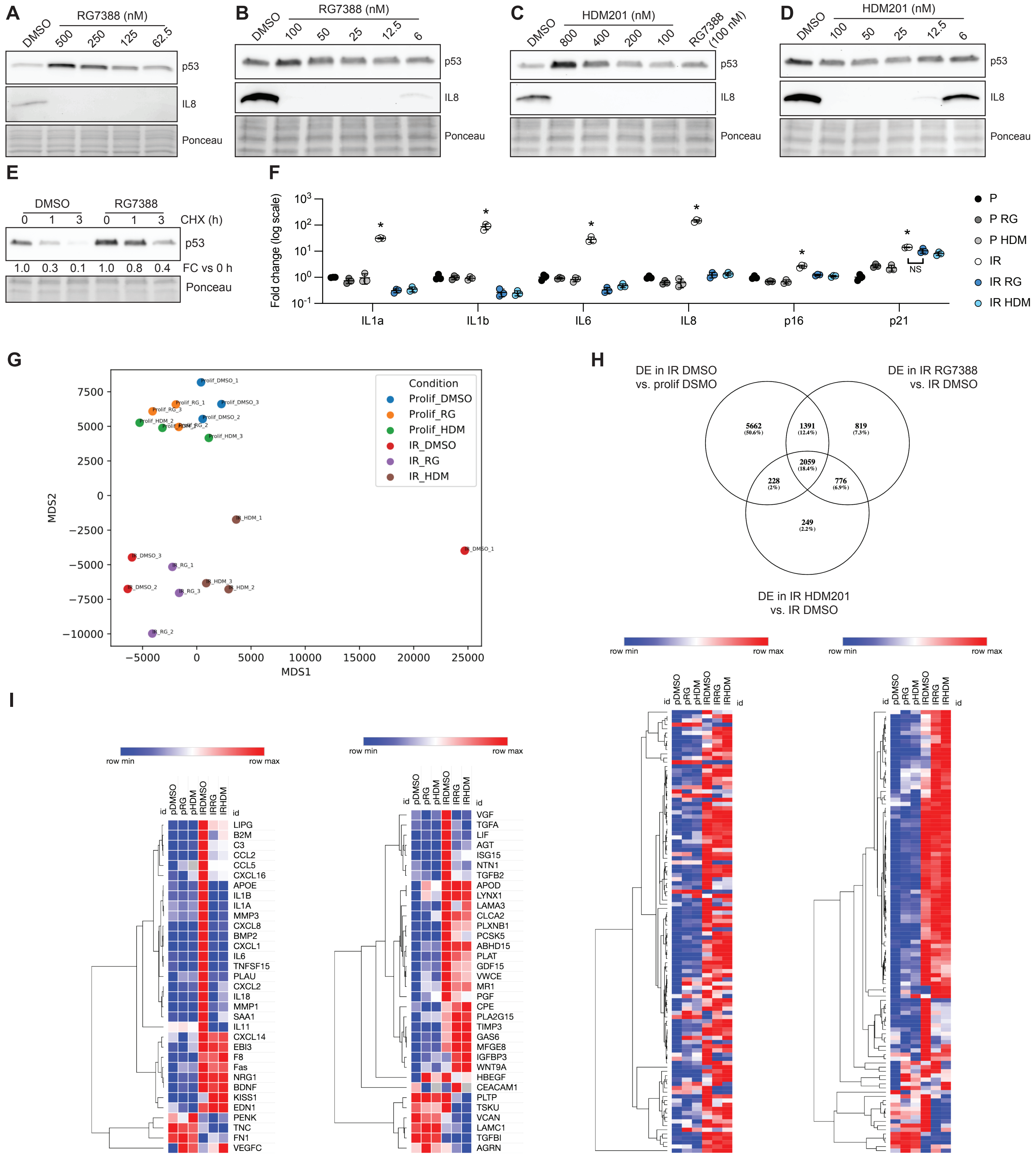


**Figure S2: Related to Figure 1.** A-D) MDM2i dose-response curves in irradiation-induced senescent IMR90 human fibroblasts by IL8 WB, n=1 experiment each. E) Cycloheximde chase assay in IMR90 cells 4 days after irradiation, measuring p53 protein level by WB and quantitation as fold change vs. 0 h for each group, representative of n=2 experiments. F) qPCR validation and G) RNA-seq principal component analysis for MDM2i treatment related to Fig.1G. H) Venn diagram of DE genes. I) Left to right: NFkB, p53, p21, and p16-associated secretomes. Data shown as means ±SD, asterisk(*) indicates p<0.05 by Student's t-test. P: proliferating control; IR: ionizing irradiation-induced senescence; CHX: cycloheximide.

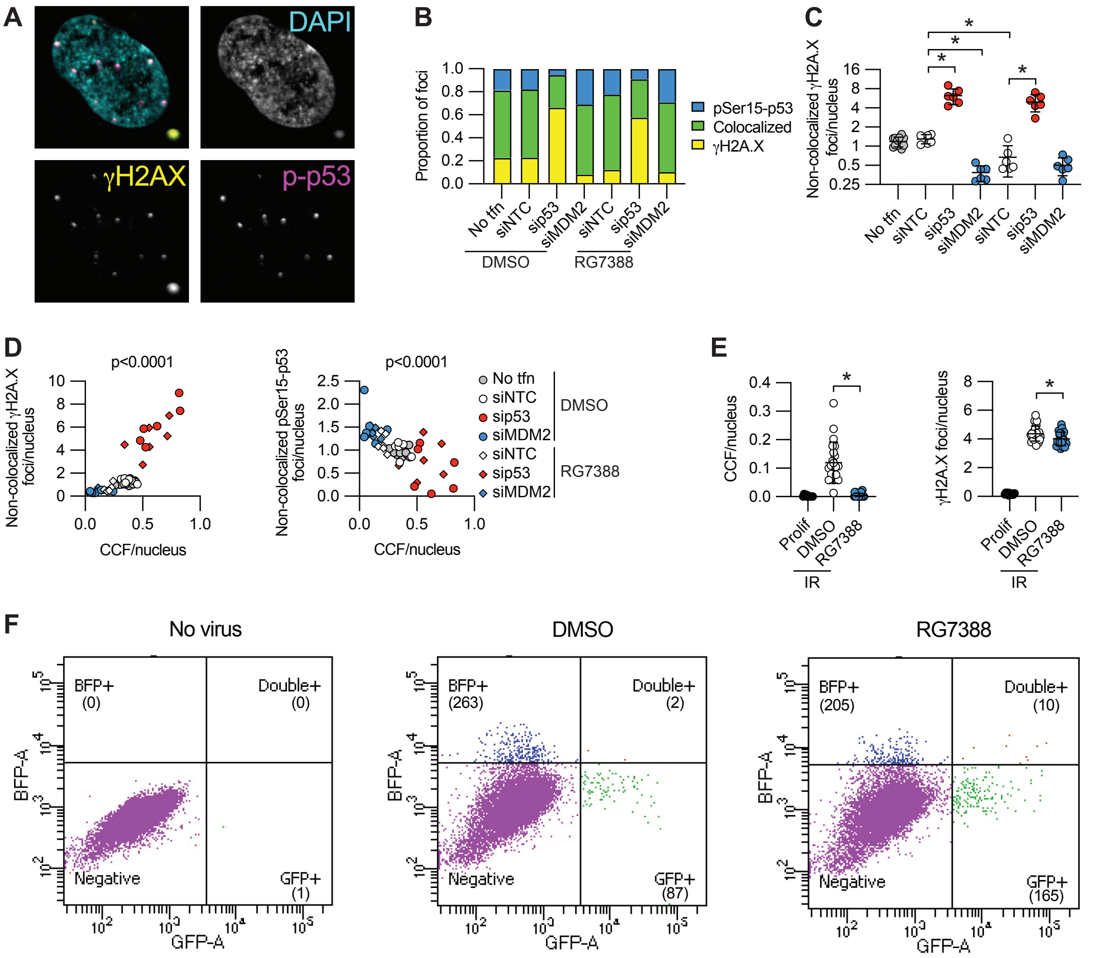


**Figure S3: Related to Figure 2.** A) Representative IF image of γH2A.X and pS15-p53 colocalization in irradiation-induced senescent IMR90 human fibroblasts and B) IF quantitation of colocalization between pS15-p53 and γH2A.X, representative of n=2 experiments. C) Quantitation of γH2A.X foci not colocalized with pS15-p53, and D) correlations to CCF formation of (left) γH2A.X foci not colocalized with pS15-p53 foci and (right) pS15-p53 foci not colocalized with γH2A.X foci, from same dataset as Fig.1B. E) Irradiation-induced senescent I9A human fibroblasts, showing CCF formation and nuclear γH2A.X foci by IF, representative of n=2 experiments. F) Representative raw data related to Fig.2G. Data shown as means ±SD, asterisk(*) indicates $p<0.05$ by Student's t-test. Prolif: proliferating control; NTC: non-targeting control.

**A** Run 1 (LIANTI)

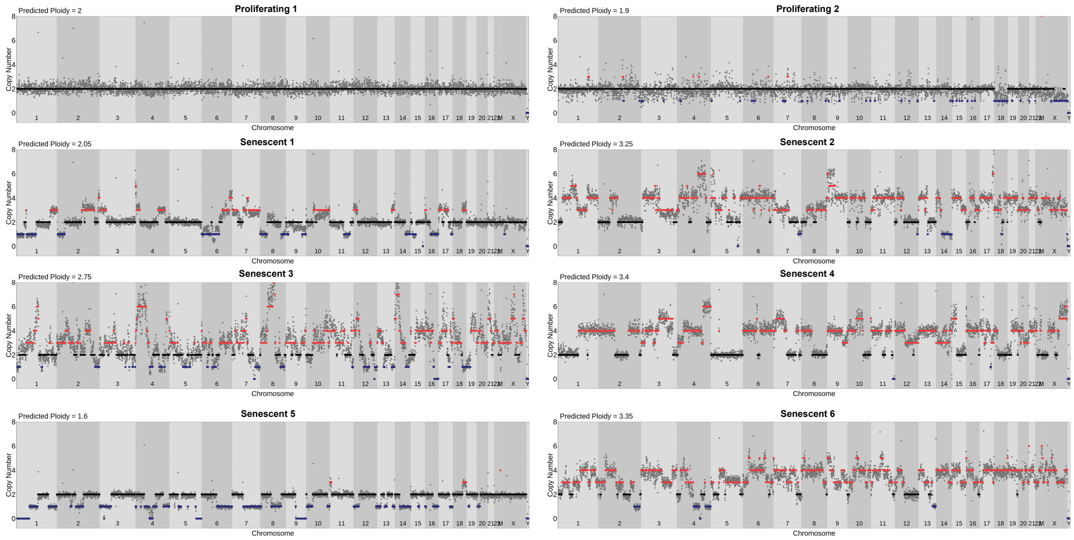


**B** Run 2 (PicoPLEX Gold)

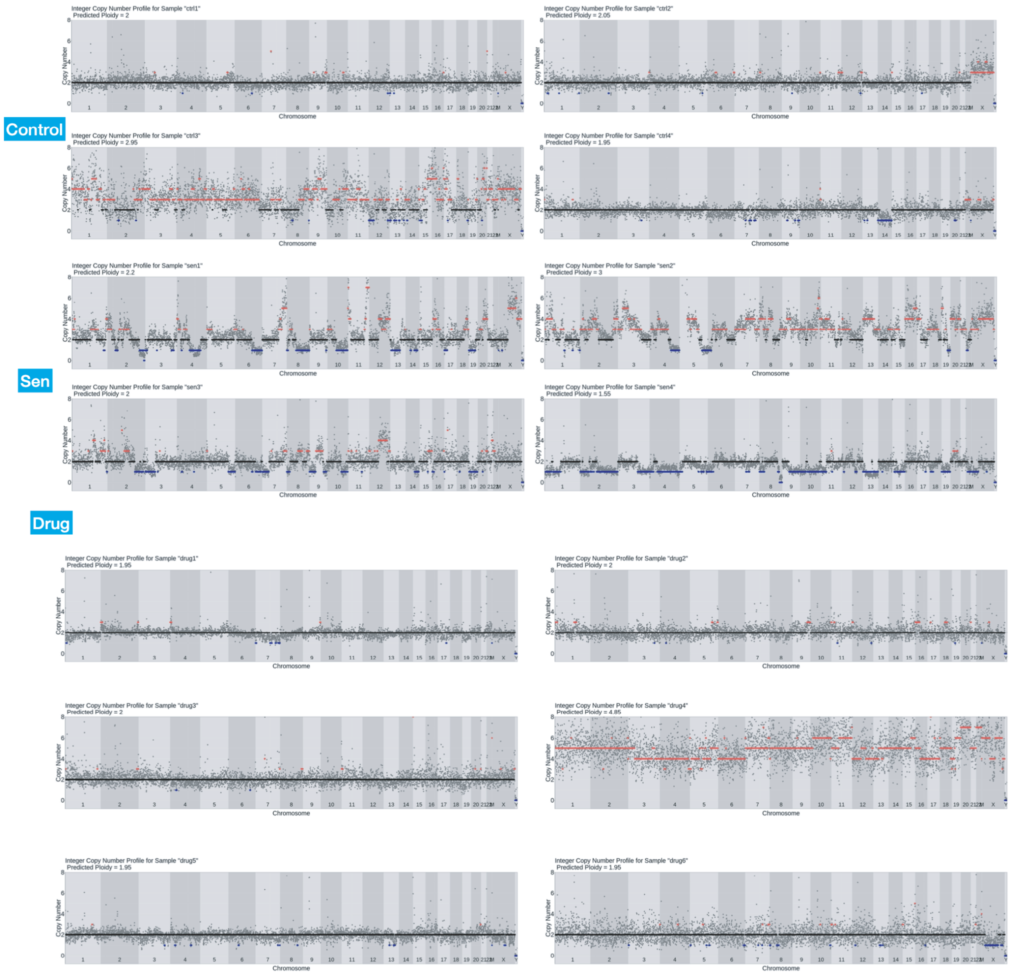


**Figure S4: Related to Figure 3.** Whole genome copy number variation plots for all cells sequenced by A) LIANTI or B) PicoPLEX Gold approaches.

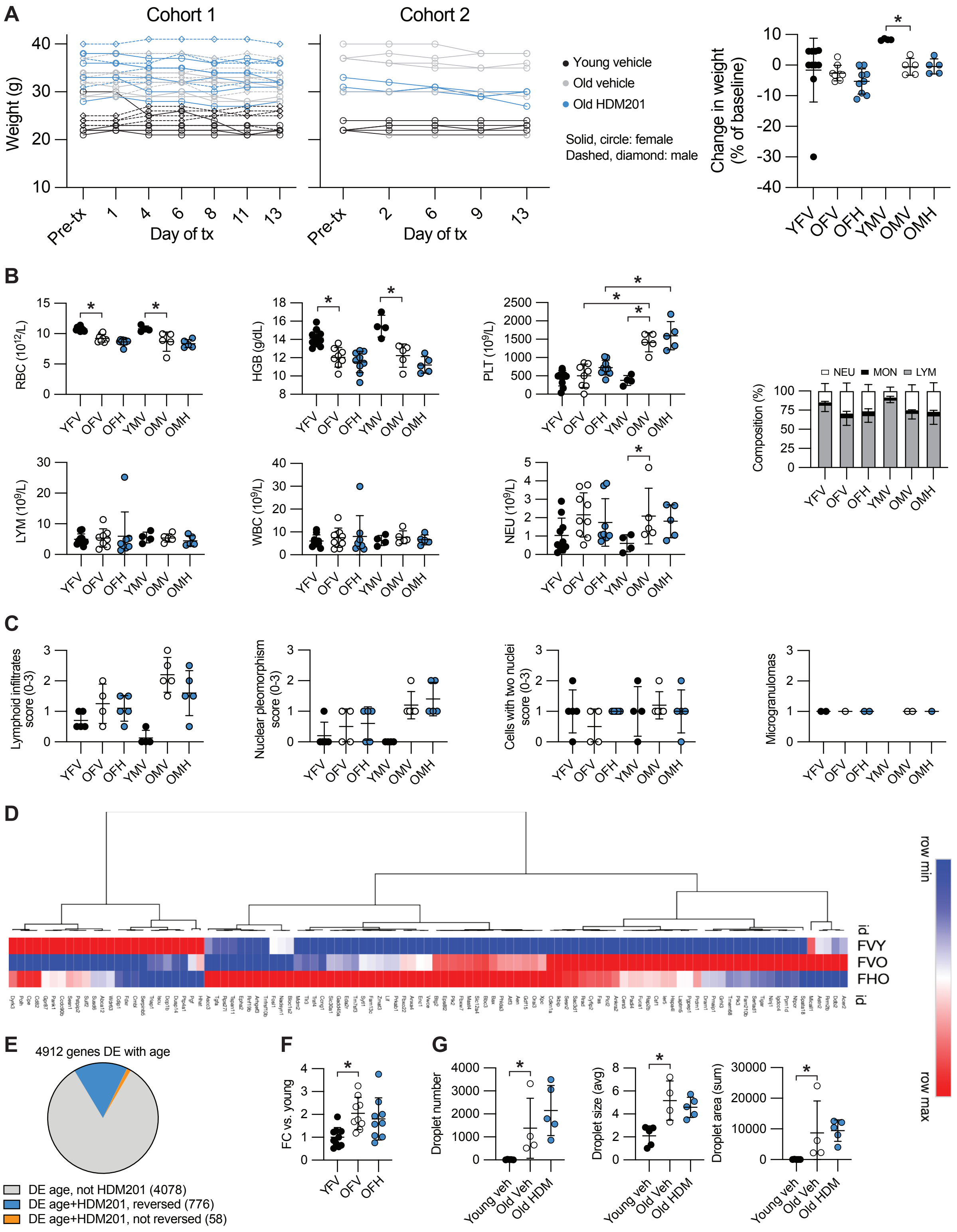

**Figure S5: Related to Figure 4.** A) Body weight and change in weight over the course of treatment as a percentage of baseline animal weight, n=4-10 per group. B) Whole blood measures and white blood cell composition, n=4-10. C) Liver pathology indices scored by a trained pathologist, n=4-10. D) p53 target gene expression in female mouse liver by RNA-seq, n=4-5 per group. E) Comparison of genes differentially expressed with age and HDM201 in liver by bulk RNA-seq, n=4-5 per group. F) Picosirius red staining in female mice, n=9-10. G) Oil red-O staining, n=4-5. Data shown as means ±SD, asterisk(*) indicates p<0.05 by Mann-Whitney U test. V: vehicle; H: HDM201; Y: young; O: old; F: female; M: male.

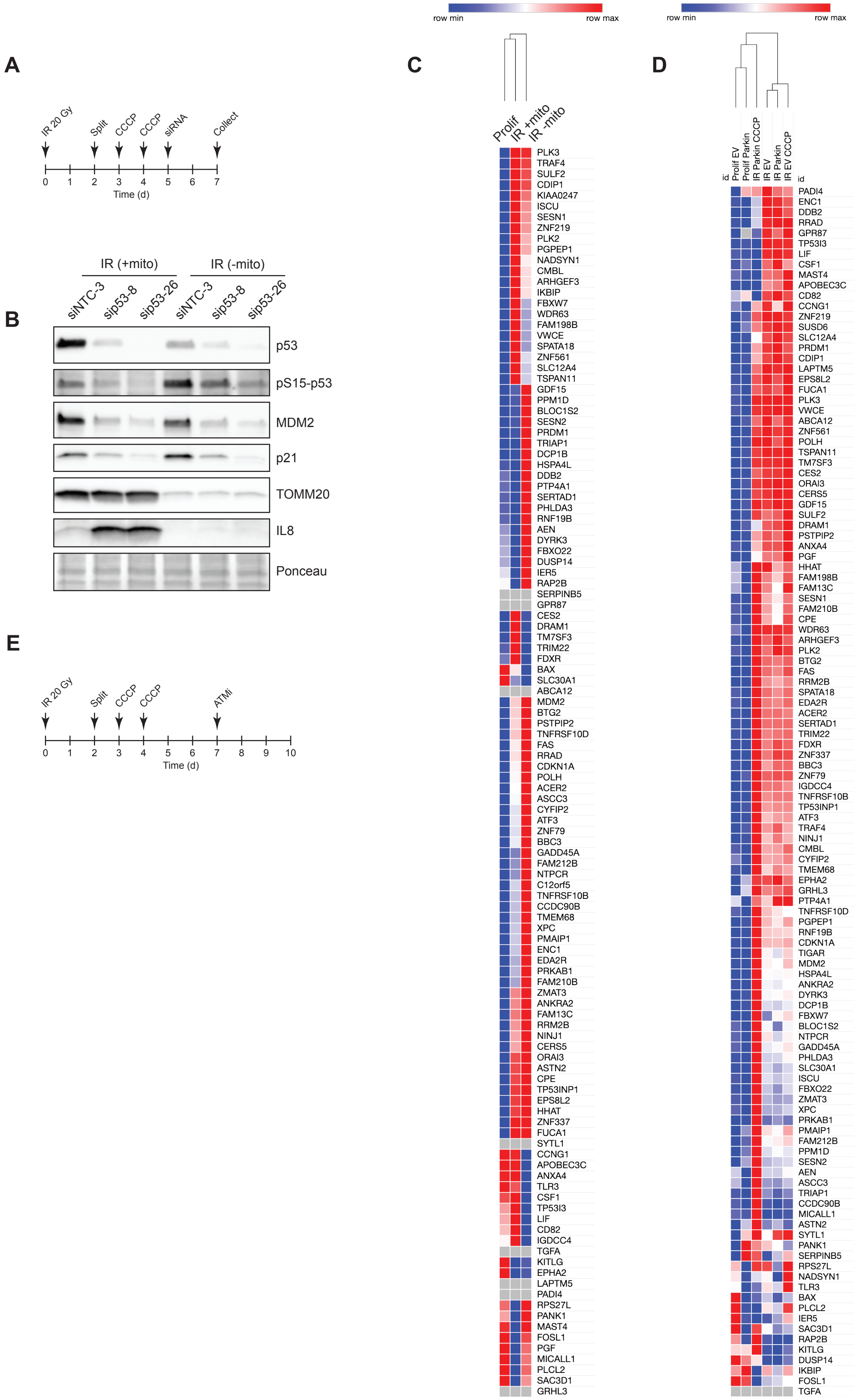


**Figure S6: Related to Figure 5.** A) Timeline for mitochondrial ablation experiments in Fig.5 A,B. B) WB validation of mitochondrial ablation and p53 knockdown related to Fig.5A,B. C) p53 target genes with mitochondrial ablation in MRC5 fibroblasts, from ref[51]. D) Heatmap related to Fig.5C, showing additional controls. E) Timeline for mitochondrial ablation experiment with ATMi treatment in Fig.5D. IR: ionizing radiation-induced senescence.

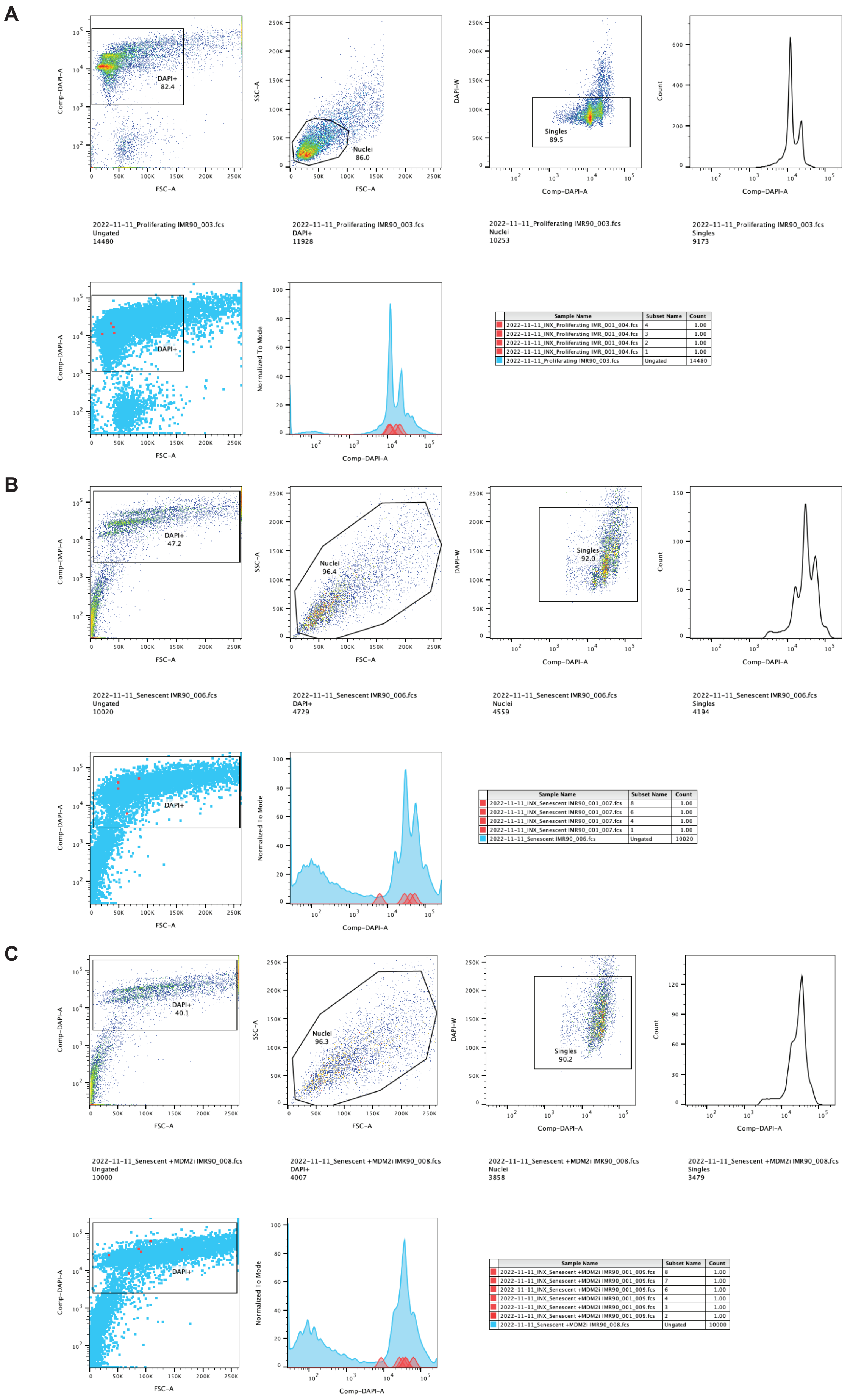


**Figure S7: Gating of cell sorting for single-nucleus genome resequencing.** A) Proliferating, B) Senescent, C) Senescent + MDM2i. Red marks correspond to the individual sorted nuclei from Fig.S4B.

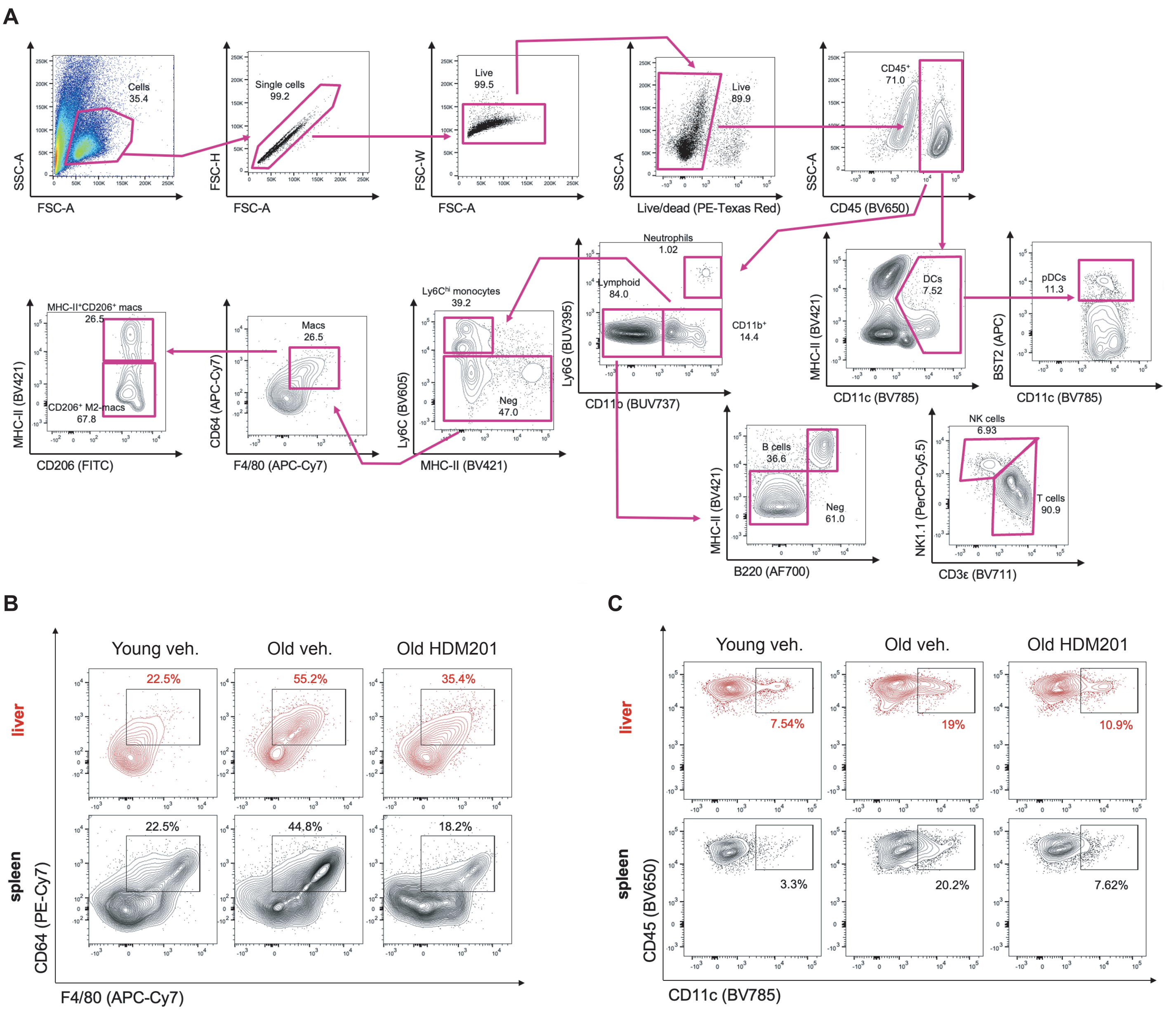


**Figure S8: Flow cytometry gating for immune profiling related to Figure 4.** A) Representative gating strategy for identifying immune cells mouse liver. B) Representative flow cytometry plots showing the frequency of macrophages (F4/80+CD64+) and C) dendritic cells (CD11c+) isolated from liver and spleen. The plots in B and C are gated on CD45+CD11b+Ly6C- cells.
